# A simulation framework for modeling the within-patient evolutionary dynamics of SARS-CoV-2

**DOI:** 10.1101/2023.07.13.548462

**Authors:** John W Terbot, Brandon S. Cooper, Jeffrey M. Good, Jeffrey D. Jensen

**Affiliations:** Arizona State University, School of Life Sciences, Center for Evolution & Medicine, Tempe, Arizona, United States of America; University of Montana, Division of Biological Sciences, Missoula, Montana, United States of America

**Keywords:** evolutionary genomics, population genetics, SARS-CoV-2, viral evolution

## Abstract

The global impact of Severe Acute Respiratory Syndrome Coronavirus 2 (SARS-CoV-2) has led to considerable interest in detecting novel beneficial mutations and other genomic changes that may signal the development of variants of concern (VOCs). The ability to accurately detect these changes within individual patient samples is important in enabling early detection of VOCs.

Such genomic scans for positive selection are best performed via comparison of empirical data to simulated data wherein evolutionary factors, including mutation and recombination rates, reproductive and infection dynamics, and purifying and background selection, can be carefully accounted for and parameterized. While there has been work to quantify these factors in SARS-CoV-2, they have yet to be integrated into a baseline model describing intra-host evolutionary dynamics. To construct such a baseline model, we develop a simulation framework that enables one to establish expectations for underlying levels and patterns of patient-level variation. By varying eight key parameters, we evaluated 12,096 different model-parameter combinations and compared them to existing empirical data. Of these, 592 models (∼5%) were plausible based on the resulting mean expected number of segregating variants. These plausible models shared several commonalities shedding light on intra-host SARS-CoV-2 evolutionary dynamics: severe infection bottlenecks, low levels of reproductive skew, and a distribution of fitness effects skewed towards strongly deleterious mutations. We also describe important areas of model uncertainty and highlight additional sequence data that may help to further refine a baseline model. This study lays the groundwork for the improved analysis of existing and future SARS-CoV-2 within-patient data.

**Significance Statement:** Despite its tremendous impact on human health, a comprehensive evolutionary baseline model has yet to be developed for studying the within-host population genomics of SARS-CoV-2. Importantly, such modeling would enable improved analysis and provide insights into the key evolutionary dynamics governing SARS-CoV-2 evolution. Given this need, we have here quantified a set of plausible baseline models via large-scale simulation. The commonly shared features of these relevant models - including severe infection bottlenecks, low levels of progeny skew, and a high rate of strongly deleterious mutations - lay the foundation for sophisticated analyses of SARS-CoV-2 evolution within patients using these baseline models.

## Introduction

The emergence of Severe Acute Respiratory Syndrome Coronavirus 2 (SARS-CoV-2) in late 2019 is the most impactful human pathogen to arise thus far in the 21^st^ century. Since its emergence, SARS-CoV-2 is directly responsible for nearly 8 million deaths as of December 2022 (Institute for Health Metrics and Evaluation 2022, Hay et al. 2023). However, the true impact of SARS-CoV-2—considering under-reporting, late reporting, and indirect deaths (*e.g.*, via strained health care systems)— is likely much greater, with worldwide excess mortality estimated to exceed 14 million as of December 2021 (Wang et al. 2022, Msemburi et al. 2023). Moreover, SARS-CoV-2 continues to persist worldwide and appears likely to become an endemic virus going forward, as observed with other human coronaviruses (HCoV-229E, -NL63, -OC43, and -HKU1; Corman et al. 2018).

Understanding the evolutionary dynamics of SARS-CoV-2 and predicting new Variants of Concern (VOCs) that may result in waves of increased infection and mortality remains vital. While the tools for studying these questions as framed via inter-host spread of SARS-CoV-2 have been well-developed (Rambaut et al. 2020, O’Toole et al. 2021), the ability to study intra-host evolution of SARS-CoV-2 is considerably less established. Transmission between hosts is an obviously important stage in viral spread; however, inter-host spread is a brief portion of the viral life cycle, with the entirety of viral reproductive activity occurring within a host. Thus, dissecting the intra-host evolutionary dynamics are of key importance in monitoring contemporary and future SARS-CoV-2 spread. In particular, mutations that influence evasion of the host immune system, increase success in cell invasion, and otherwise improve the successful completion of metabolic and reproductive tasks within a host cell could all be of clinical consequence. Given complete information, such mutations would first be detectable within a single host. While strongly beneficial mutations can eventually become identifiable when observing inter-host data through their increased prevalence within the meta-population (should they escape stochastic loss), the evolutionary dynamics that ultimately dictate their spread will be determined at the intra-host level.

For the viral population within a single patient, these episodic, beneficial mutations may be expected to modify patterns of genomic variation in a manner that would deviate from background patterns produced under constantly operating evolutionary processes (*e.g.*, via a selective sweep of the beneficial mutation (Charlesworth and Jensen 2021). As such, as with any natural population, the study of viral intra-host evolution requires the construction of an evolutionary ‘null’ model to quantify expected baseline levels and patterns of genomic variation (Jensen 2021, Johri et al. 2020, Terbot et al. 2023). At a minimum, a viral baseline model should include mutation, recombination, reproductive dynamics, purifying and background selection, and the history of bottlenecks and growth characterizing patient infection (Irwin et al. 2016). Without comparison to such a baseline model, it is not possible to determine if observed within-patient allele frequencies are attributable to these common evolutionary processes or to the comparatively rare action of positive selection (Barton 1998, Johri et al. 2022b).

In this study, we present a first attempt to construct such a model and narrow the range of key parameter values governing the intra-host evolutionary dynamics of SARS-CoV-2. Using forward simulations and comparison to existing patient data summarizing intra-host variation, we identify more and less likely areas of parameter space. We find that transmission and infection bottlenecks appear to be strongly constrained (*i.e.,* on the order of <5 virions) in plausible models, consistent with other recent results for SARS-CoV-2 (Lythgoe et al. 2021, Martin and Koelle 2021, Bendall et al. 2023) and the transmission of other airborne viruses like seasonal influenza (McCrone et al. 2018, Valesano et al. 2019). We also describe important areas of uncertainty and correlations between parameter values. For example, if the distribution of new mutational effects is heavily skewed towards strongly deleterious mutations, the range of uncertainty in other parameter values is inflated. Furthermore, we highlight additional data that may help to further narrow this parameter space. For example, the commonly employed 2% minor allele frequency threshold cut-off greatly limits model resolution, and may be improved by higher-coverage sequencing of individual patient samples that increases the confidence in individual single nucleotide polymorphism (SNP) calls. Thus, the presented exploration of the SARS-CoV-2 evolutionary parameter space will be valuable in informing future modeling studies, in interpreting newly emerging patient data, and in guiding future data collection.

## Results

A total of 12096 model-parameter combinations were analyzed (Table 1, Table 2, and Figure 1). The great majority were rejected for generating too few or too many segregating SNPs relative to available empirical data, leaving 592 models remaining (Supplemental Table 1). Due to the nested nature of the models, the feasible model-parameter sets can be readily categorized according to the 108 possible combinations of the required parameters. Of these, 18 specific parameter combinations produced viable models, highlighting which parameters are likely to have the strongest influence of intra-host evolution of SARS-CoV-2. We found that all plausible models included a strong infection bottleneck *(i.e*., a bottleneck under 5 virions appears most consistent with the data, represented here by a bottleneck of 1; Figure 2, Figure 3). Most retained models also had low or moderate mutation rates. Only three models with the highest mutation rate were retained. All three of these models were full models using a distribution of fitness effects (DFE) with the largest proportion of strongly deleterious mutations, the highest carrying capacity, and briefest infection duration; the parameter values for recombination and progeny skew varied between these models. In contrast, all 12 required parameter sets using the lowest mutation rate and a severe bottleneck of 1 cleared the SNP threshold, and 5 of the 12 sets with the midpoint mutation rate resulted in plausible models. Generally, larger carrying capacities resulted in better fitting models (4/12 parameter sets had plausible models with the lowest value of carrying capacity and a bottleneck of 1, 6/12 with the midpoint value for carrying capacity, and 8/12 for the highest carrying capacity).

**Figure 1.**
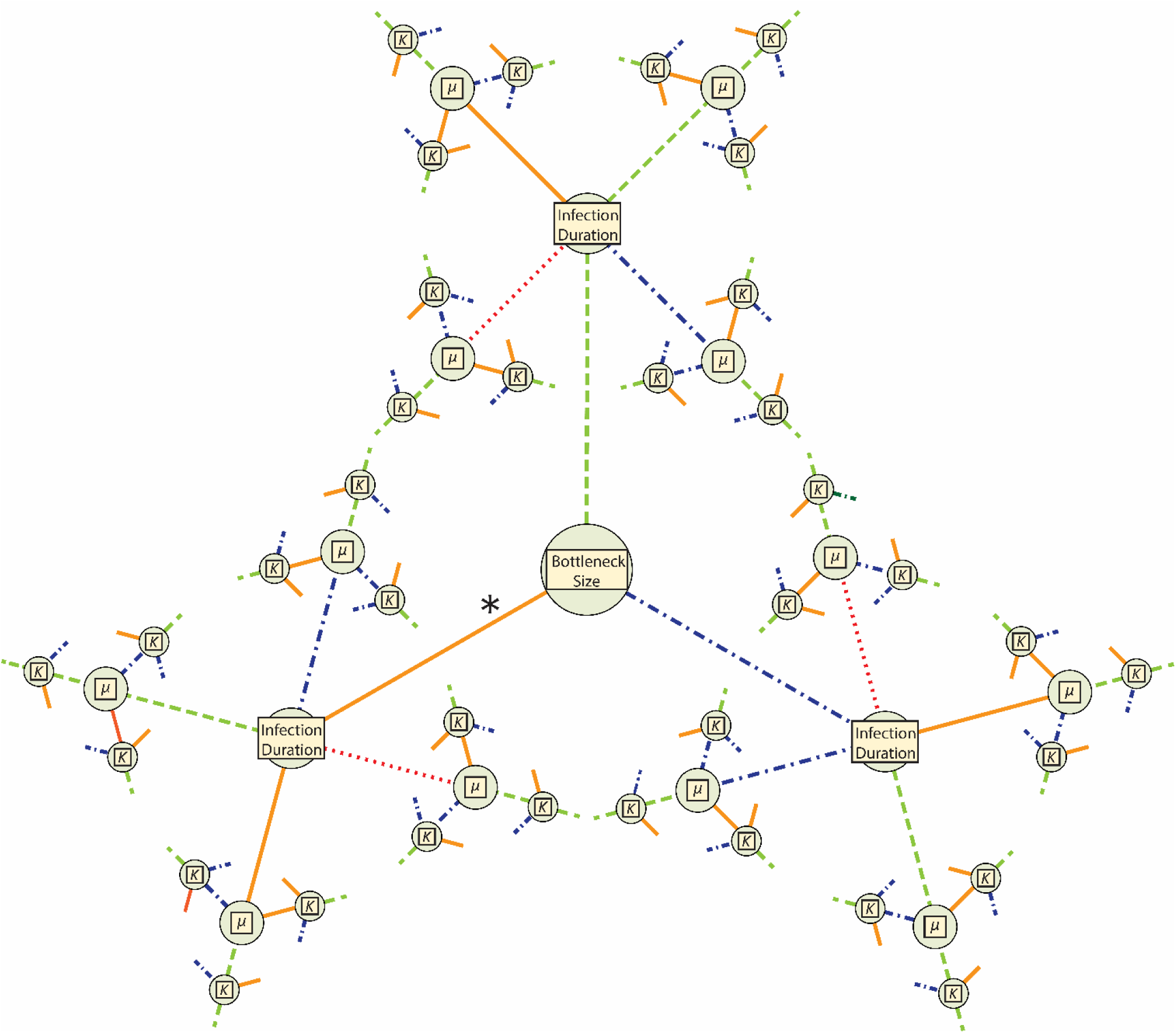
Representations of the total parameter space for models’ required parameters. Each required parameter [*i.e.*, bottleneck size, infection duration, mutation rate (*μ*), and carrying capacity (*K*)] is represented by a node. Line color and shape correspond to parameter value: dotted, red is lowest; solid, orange is low (the lowest value for bottleneck size, *μ*, and *K*); dashed, green is the mid-point value; and dotted-dashed, blue is the high value. The asterisk (*) denotes the only area of parameter space that had plausible models: those with bottlenecks of 1 (see Figure 2).

**Figure 2.**
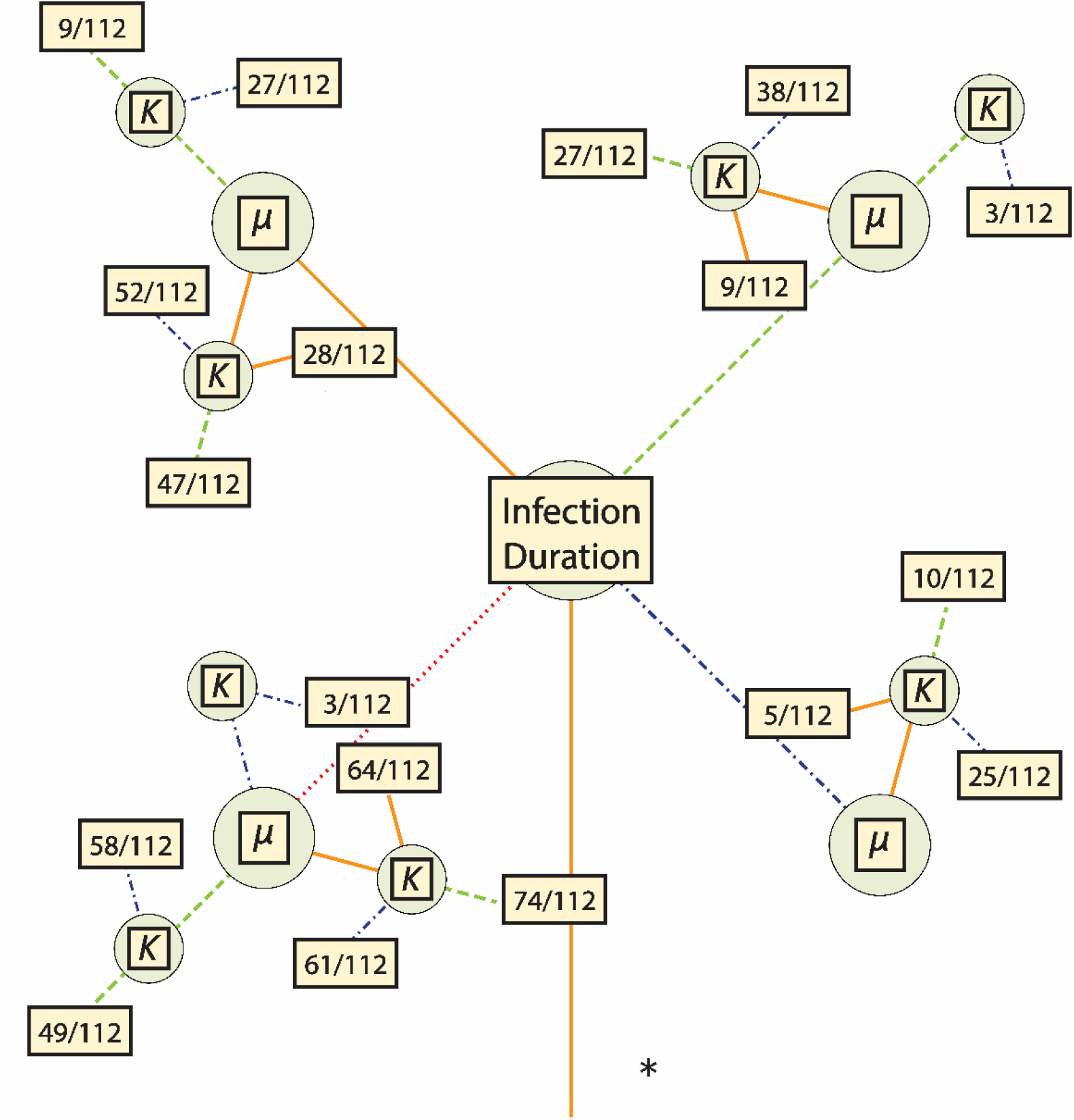
Parameter space for all plausible models. Line color and shape correspond to the parameter values as in Figure 1. For each set of required parameters, a box is included that contains the fraction of plausible models out of the total number of models using that set. Further information on the specific models which were plausible is detailed in Table 3.

**Figure 3.**
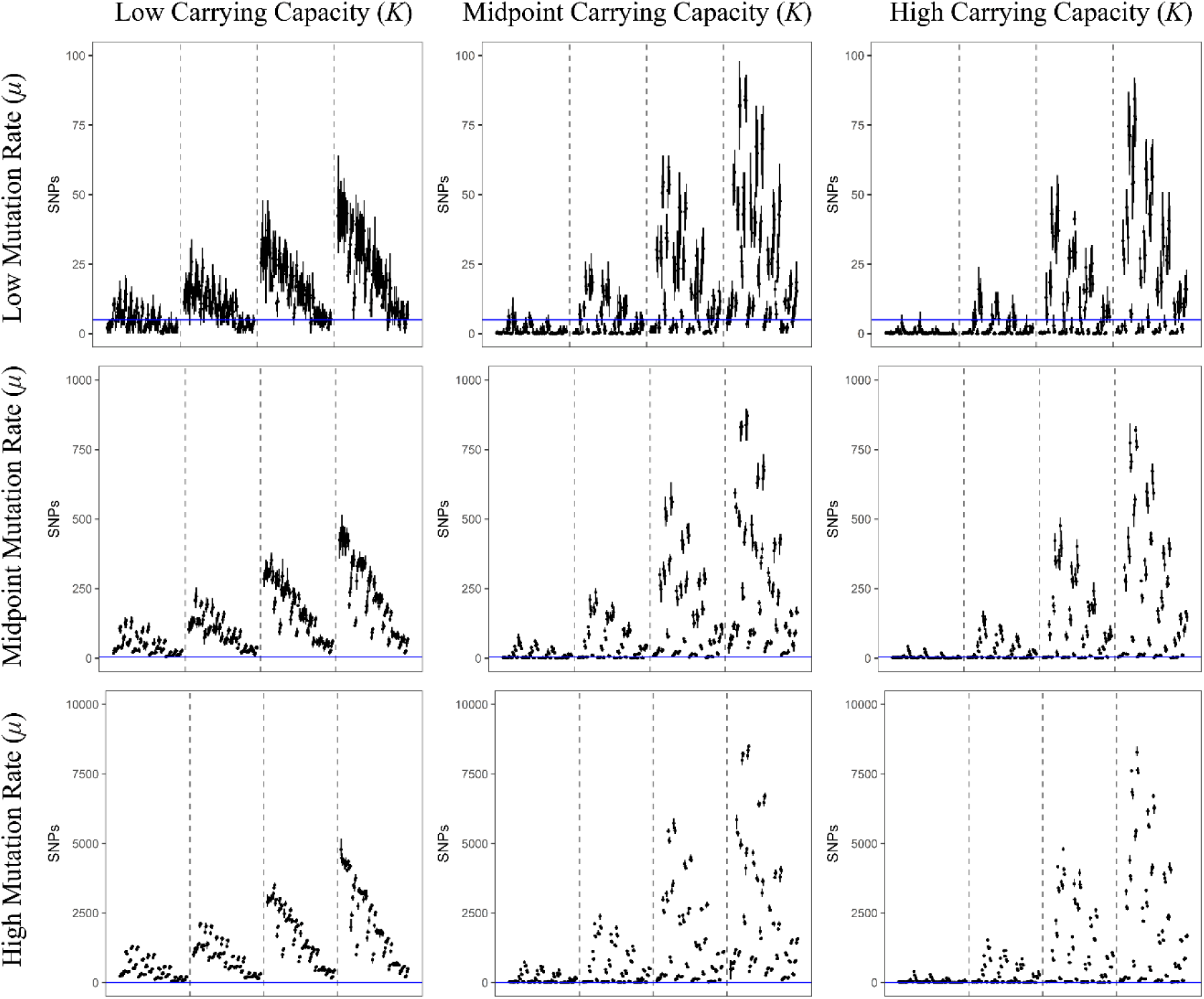
Each line represents the range of filtered SNPs for a particular model using a sampling of 1000 genomes, and each point is the mean of that model’s replicates. All models in this figure used the lowest bottleneck size (*i.e.*, 1); the carrying capacity used increases in panels from left to right, and the value of the mutation rate used increases in panels from top to bottom. Within each panel, dashed lines separate models into subpanels with different infection durations, increasing from left to right. Within each subpanel the order of the models is the same and is detailed in Supplemental Table 3. The blue, horizontal line represents the threshold of 5 SNPs used in this study to accept or reject a potential model (note that Y-axes differ by row). Models with a mean value lower than this line (but not zero) were accepted as plausible. Similar figures for models using larger bottlenecks are available as Supplemental Figures 1 and 2.

**Table 1.** Outline of Model Complexity Each row represents a nested model which includes all parameters in that row and from rows above it. Number of parameters added by each stage is detailed in column “+*k*” and the specific parameters added are described in the rightmost column.

| Model | +$k$ | Parameters |
| --- | --- | --- |
| Bottleneck | 4 | Bottleneck, Mutation Rate ($\mu$), Infection Duration, Carrying Capacity ($K$) |
| + Recombination | 1 | Recombination Rate ($R$) |
| + Progeny Skew | 2 | Probability of Multiple Coalescent Event ($\Xi$), Burst Size |
| + DFE | 1 | Ratio of neutral to strongly deleterious mutations |

**Table 2.** Parameter Values Used in Models Each parameter, other than infection duration, had three possible levels: a low point estimate, a high point estimate, and a value representing a midpoint value. The column labeled “Burn-In” details the parameter levels used during the common burn-in; note that aside from infection duration and carrying capacity, all parameter levels used were the midpoint level. The highest carrying capacity and an extended infection duration were used to allow mutations to accumulate and begin to reach equilibrium as may be expected across the entire metapopulation of a pathogen during a prolonged pandemic.

|  | Burn-In | Lowest | Low | Midpoint | High |
| --- | --- | --- | --- | --- | --- |
| Initial Bottleneck | N/A | N/A | 1 | 5 | 100 |
| Infection Duration | 25e3 | 168 (7 days) | 336 (14 days) | 672 (28 Days) | 1008 (42 Days) |
| Mutation Rate | 2.135e-6 | N/A | 2.135e-7 | 2.135e-6 | 2.135e-5 |
| Carrying Capacity | 1e5 | N/A | 5e3 | 5e4 | 1e5 |
| Recombination Rate | 5.5e-5 | N/A | 1e-5 | 5.5e-5 | 1e-4 |
| Progeny Skew Probability | 0.003 | N/A | 0.0001 | 0.003 | 0.1 |
| Progeny Skew Amount | 100 | N/A | 20 | 100 | 200 |
| DFE (Neutral:Deleterious) | 1:1 | N/A | 4:1 | 1:1 | 1:4 |

**Table 3:** Number of Plausible Models Given Required Parameters and Model Complexity Counts and percentage of nested models within a given set of values for the required parameters carrying capacity, mutation rate, and infection duration detailed in the first three columns (the initial bottleneck for all accepted models was the low value or a single virion). Each set of required parameters had a single bottleneck model (columns 4 and 5), 3 recombination models (columns 6 and 7), 27 progeny skew models (columns 8 and 9), and 81 full models (columns 10 and 11). The final two columns represent the count and percentage of accepted models (out of a possible 112) given a set of required parameters.

| Infection Duration | Mutation Rate ($\mu$) | Carrying Capacity ($K$) | Bottleneck Count | Bottleneck Percent | Recomb. Count | Recomb. Percent | Prog. Skew Count | Prog. Skew Percent | Full Count | Full Percent | All Count | All Percent |
| --- | --- | --- | --- | --- | --- | --- | --- | --- | --- | --- | --- | --- |
| Lowest | Low | Low | 1 | 100.00% | 2 | 66.70% | 6 | 22.20% | 55 | 67.90% | 64 | 57.10% |
| Lowest | Low | Midpoint | 1 | 100.00% | 3 | 100.00% | 14 | 51.90% | 56 | 69.10% | 74 | 66.10% |
| Lowest | Low | High | 1 | 100.00% | 2 | 66.70% | 19 | 70.40% | 39 | 48.10% | 61 | 54.50% |
| Lowest | Midpoint | Midpoint | 1 | 100.00% | 3 | 100.00% | 8 | 29.60% | 37 | 45.70% | 49 | 43.80% |
| Lowest | Midpoint | High | 1 | 100.00% | 3 | 100.00% | 11 | 40.70% | 43 | 53.10% | 58 | 51.80% |
| Lowest | High | High | 0 | 0.00% | 0 | 0.00% | 0 | 0.00% | 3 | 3.70% | 3 | 2.70% |
| Low | Low | Low | 0 | 0.00% | 0 | 0.00% | 0 | 0.00% | 28 | 34.60% | 28 | 25.00% |
| Low | Low | Midpoint | 1 | 100.00% | 3 | 100.00% | 10 | 37.00% | 33 | 40.70% | 47 | 42.00% |
| Low | Low | High | 0 | 0.00% | 3 | 100.00% | 12 | 44.40% | 40 | 49.40% | 52 | 46.40% |
| Low | Midpoint | Midpoint | 0 | 0.00% | 0 | 0.00% | 0 | 0.00% | 9 | 11.10% | 9 | 8.00% |
| Low | Midpoint | High | 0 | 0.00% | 3 | 100.00% | 3 | 11.10% | 21 | 25.90% | 27 | 24.10% |
| Midpoint | Low | Low | 0 | 0.00% | 0 | 0.00% | 0 | 0.00% | 9 | 11.10% | 9 | 8.00% |
| Midpoint | Low | Midpoint | 1 | 100.00% | 2 | 66.70% | 1 | 3.70% | 23 | 28.40% | 27 | 24.10% |
| Midpoint | Low | High | 1 | 100.00% | 3 | 100.00% | 9 | 33.30% | 25 | 30.90% | 38 | 33.90% |
| Midpoint | Midpoint | High | 0 | 0.00% | 0 | 0.00% | 0 | 0.00% | 3 | 3.70% | 3 | 2.70% |
| High | Low | Low | 0 | 0.00% | 0 | 0.00% | 0 | 0.00% | 5 | 6.20% | 5 | 4.50% |
| High | Low | Midpoint | 0 | 0.00% | 0 | 0.00% | 0 | 0.00% | 10 | 12.30% | 10 | 8.90% |
| High | Low | High | 1 | 100.00% | 1 | 33.30% | 0 | 0.00% | 23 | 28.40% | 25 | 22.30% |

Within these required parameter sets, there are a total of 112 nested models (1 bottleneck model, 3 with the addition of recombination, 27 with the additions of recombination and progeny skew, and 81 with the additions of recombination, progeny skew, and a DFE); the number of models that cleared the filtering requirement varied considerably between the required parameter sets. Briefly, 9 of the 18 required parameter sets had bottleneck models consistent with the empirical data (of which 8 had examples of all nested models also clear the threshold), 11 of the 18 had consistent recombination-only models, 10 out of 18 had consistent recombination + progeny skew models, and all 18 had consistent full models. Generally, if a simpler model could be accepted, then more complicated models nested within that model also tended to produce plausible results. The distribution of these non-rejected models are more fully detailed in Table 3 and Figure 2. In terms of general patterns, models with strong infection bottlenecks, lower progeny skews, and DFEs containing a greater proportion of strongly deleterious mutations were more likely to be accepted (Figure 3, and see Discussion).

## Discussion

The evaluation of intra-host population genomic data from SARS-CoV-2 patient samples has the potential to detect signatures of positive selection and allow for the early identification of VOCs. However, it is important to recognize that levels and patterns of genetic variation in a population are the result of a variety of factors including mutation, recombination, reproductive dynamics, purifying and background selection, and the history of bottlenecks and growth characterizing patient infection. One of the key methods of differentiating between these alternative explanations is through comparisons between empirical data and plausible simulated data (Irwin et al. 2016, Johri et al. 2020, Jensen 2021). However, given that the possible model and parameter space is essentially infinite, a key first step in developing an evolutionary baseline model is determining the relevant parameter bounds.

Our study supports three main conclusions that help define a feasible parameter space governing intra-host SARS-CoV-2 evolutionary dynamics. First, our results support recent conclusions that SARS-CoV-2 transmission is generally associated with strong infection bottlenecks of one or a few virions (Bendall et al. 2023). Specifically, we found a bottleneck of 1 to be the only bottleneck size tested that produced plausible models. However, the unexamined space between 1 and 5 (the midpoint value for bottlenecks) may also produce plausible models as they were not tested in this study. Regardless, a bottleneck of 1 to 4 virions conforms with the recent empirical findings based on patterns of intra-host SARS-CoV-2 variation between likely transmissions pairs (Bendall et al. 2023).

Secondly, our results highlight the importance of considering the pervasive effects of purifying and background selection, via a consideration of the DFE, in constraining levels of variation segregating above the 2% frequency threshold. The wide-spread production of strongly deleterious mutations appears necessary to explain the apparent contradiction between the high mutation rates of RNA viruses (Drake and Holland 1999, Elena and Sanjuán 2005, Jensen et al. 2020) and the generally low number of SNPs identified in patients ((Lythgoe et al. 2021, Valesano et al. 2021, Wang et al. 2021, Bendall et al. 2023, Gu et al. 2023). Given that coding regions comprise most of the SARS-CoV-2 genome, this result is not particularly surprising. This importance may be observed in the simulated parameter sets: among the models that accumulated only neutral mutations, 11.8% were plausible; among the models with primarily neutral mutations, 23.9% were plausible; among the models with equivalent rates of neutral and strongly deleterious mutations, 28.4% were plausible; and among the models with primarily strongly deleterious mutations, 42.8% were plausible. Current best estimates of the true DFE underlying SARS-CoV-2 intra-host evolution suggest a bimodal distribution of fitness effects (Flynn et al. 2022, Terbot et al. 2023) with peaks centered around strongly deleterious and neutral fitness impacts. At a minimum, between 15-20% of sites in the SARS-CoV-2 genome seem to be entirely invariant (Neher 2022), providing a minimum estimate for the proportion of strongly deleterious mutations in the DFE. However, other studies indicate this value is more likely in the range of 40-50% (Flynn et al. 2022, Terbot et al. 2023). Therefore, while the DFE most severely skewed towards strongly deleterious mutations produced the most plausible models, it is unlikely to reflect the true proportion of strongly deleterious mutations. However, it is notable that the two other DFE distributions reflecting current best understandings of the true DFE both produced around twice as many viable models as models including only neutral mutations, emphasizing the central importance of purifying selection in the evolution of SARS-CoV-2 within patients.

Finally, lower progeny skew values produce more plausible models. Specifically, models with progeny skew and containing either lower probabilities of skewed offspring events and/or lower burst sizes were over-represented amongst acceptances (397/555 plausible models or 71.5%). Reproduction of SARS-CoV-2 within a host cell is generally non-lytic and instead involves release of new virions through continuous budding (Bar-On et al. 2020; Park et al. 2020). The prominence of lower levels of progeny skew in plausible models is consistent with a lack of a single large burst of reproduction (*i.e*, lower values of burst size). Alternatively, it may be related to the production of sub-genomic RNA which are not packaged directly into offspring virions, but through recombination, mutations present in their sequences can be incorporated into offspring virions (*i.e.*, lower values of *Ξ*; Langsjoen et al. 2020, Gribble et al. 2021).

Further model differentiation is limited by current data available for intra-host variation of SARS-CoV-2. The 2% allele frequency cutoff used in this study mirrors the cutoff used in several previous studies (Table 4). While this criterion successfully narrowed the relevant evolutionary parameter space in our simulations, the scarce number of SNPs are insufficient to explore more sophisticated statistics related to the site frequency spectrum or linkage disequilibrium. However, new mutations arising during a typical patient infection will be rare and thus may be missed when applying a frequency-based filter. There is of course an inherent trade-off between including lower frequency SNPs and reducing the confidence in individual SNP calls (Jacot et al. 2021). This trade-off can be partially ameliorated via the generation of high-quality and higher-depth sequencing. A rule of thumb proposed by Lauring (Lauring 2020) suggests that coverage should be ten times the inverse of the variant’s frequency. So, a coverage depth of 1000 would be required to confidently detect variants at 1% of the population, a depth of 2000 for a 0.5% variant, and so on.

**Table 4.** Summary of Empirical Studies Referenced Sampling schema and results from empirical studies that reported the number of intra-host SNPs identified in patient samples. Note that all studies apart from Lythgoe et al. 2021 utilized serial sampling within at least one patient during their data collection; this did not impact their reported number of intra-host SNPs, but did allow for confirmation of SNPs as true positives.

| Paper | Time to Sample | MAF Filter | Intra-host SNPs |
| --- | --- | --- | --- |
| Lythgoe et al. 2021 | "symptomatic individuals on admission to the hospital" | 3% | 1.4 (mean) |
| Valesano et al. 2021 | -7 to 20 days post symptom onset | 2% | 1 (median); 0-2 (IQR) |
| Wang et al. 2021 | 10 to 37 days post-symptom onset (18.09 days (mean)) | 5% | 7.33 (mean); 1-23 (Range) |
| Bendall et al. 2023 | Symptoms and positive test within 7 days | 2%* | ~0.82 (mean); 0-5 (Range) |
| Gu et al. 2023 | With 5 days of symptom onset (2.3 days (mean)) | 2.50% | 10.05 (mean); 5 (median) |

\* Intra-host SNP had to clear MAF filter in two technical replicates

These general bioinformatic recommendations assume that sequencing data will effectively sample intra-host variation without bias. However, most SARS-CoV-2 sequencing protocols rely on targeted PCR or probe-based enrichment of the SARS-CoV-2 genome to reduce background contamination of host nucleotides. Targeted enrichment approaches have been widely used in biology and are usually sensitive to any standing genetic variation that impacts the binding efficacy of enrichment probes or primers (Mamanova et al. 2010, Jones and Good 2015). Indeed, assay-dependent effects have been a constant concern during the development SARS-CoV-2 sequence protocols as demonstrated by the recurrent need to redesign enrichment primers (*e.g.*, the ARCTIC protocol) to fully sequence the genomes of newly circulating VOCs (Ulhuq et al. 2023). SARS-CoV-2 targeted enrichment is also sensitive to low viral loads (Lam et al. 2021), which could further limit understanding of changes in intra-host diversity through the time course of an infection. Given these concerns, technical replication of individual patient samples may be required to reliably detect and quantify the frequency of rare variants.

In addition to limitations relating to the detection of rare alleles, our study was also unable to consider the impacts of host compartmentalization (*i.e.*, localized subpopulations within different organs and areas of organ systems) on intra-host genetic diversity. Compartmentalization is likely an important factor for SARS-CoV-2 evolution in regard to the production of viral reservoirs in prolonged infections or infection of immune-compromised patients (Gonzalez-Reiche et al. 2023, Normandin et al. 2023). However, there is currently insufficient information regarding the number and connectivity of compartments used by SARS-CoV-2 to allow for this aspect of model complexity to be reasonably parameterized. As more empirical evidence regarding compartmentalization becomes available, future simulation studies may be able to incorporate compartmentalization to determine the impacts—specifically the influence of gene flow and recombination among compartments—on the intra-host population genetics of SARS-CoV-2.

With that said, based on the currently available data, our study has successfully quantified areas of the SARS-CoV-2 evolutionary parameter space that are most plausible. In addition, the simulation framework presented here may be utilized to compare against future sequencing studies, which will likely enable a further narrowing of likely models. However, even the relatively broad parameter space here identified may be utilized to more effectively screen for newly emerging positively selected mutations (*e.g.,* those contributing to the rapid spread of VOCs), and in so doing reduce the traditionally high false-positive rates traditionally associated with such selection scans (Johri et al 2020, Johri et al. 2022a). This work will also likely be useful in providing a baseline simulation for studies looking at intra-host population genetics of SARS-CoV-2 over time within a single host. Such studies may also allow for greater discrimination between baseline models while retaining the use of a robust, minimum minor allele frequency by providing information on how genetic variation accumulates (or persists) over the course of an infection within a single patient.

## Materials and Methods

### Simulations

We used the SLiM software package (v4.0.1, Haller and Messer 2023) to conduct forward-in-time simulations. We performed all simulations using SLiM’s non-Wright-Fisher tick cycle. To represent the single-stranded, haploid genome of one metabolically-active virion, we simulated genomes consisting of 30kb. Each tick of the simulation represented approximately one hour, during which all virions in the population produce one ‘child virion’ excluding cases of progeny skew (described further below). We ran four different, nested models to gain insights into the importance of various population genetic factors. The simplest model, hereafter referred to as the bottleneck model, consisted of four parameters: the mutation rate, the initial number of virions drawn from the burn-in period (or bottleneck size), the carrying capacity of the host, and the number of ticks over which the simulation runs *i.e.*, the time between infection and sampling (infection duration). The other three models added key factors: recombination, recombination plus progeny skew, and a full model adding recombination, progeny skew, and a DFE. The latter describes the proportion new mutations that are strongly deleterious or neutral. The model and parameter space are summarized in Table 1 and Table 2. We based the tested parameter ranges upon the current literature, as recently described in Terbot et al. 2023 unless otherwise stated.

Progeny genomes added mutations according to a single, genome-wide mutation rate. Contrary to SLiM’s default behavior for mutations arising at the same site, the simulation retained only the most recent mutation occurring in a genome. The initial bottleneck was performed by drawing virions from a common burn-in simulation, and the size of this bottleneck reflected the range reported in the literature (Popa et al. 2020, Braun et al. 2021, Lythgoe et al. 2021, Martin and Koelle 2021, Bendall et al. 2023). We based the levels used for carrying capacity on the number of virions estimated at peak infection of 10^9^ to 10^11^ (Bar-On et al. 2020, Sender et al. 2021). Due to computational constraints, and because this figure represents a census population size as opposed to an effective population size, we scaled the carrying capacity downward. Owing to this scaling of population size, we scaled up the range of mutation and recombination rates to maintain a constant population-scaled product. The duration of infection was chosen to represent a range of potential times to sampling. Given most empirical studies report sampling in terms of days from symptom onset (Table 4), and there is considerable uncertainty regarding the incubation time of SARS-CoV-2 from initial infection to symptom presentation (Bar-On et al. 2020, Lauer et al. 2020, Li et al. 2020, Du et al. 2022, Wu et al. 2022), this range extends from brief (7 days) to extended (42 days).

To model recombination, we used a single parameter for the genome-wide recombination rate. For each virion, the simulation chose another random virion to serve as the recombination partner; and the simulation used this recombination partner for all progeny produced by the focal virion in that tick cycle, including those experiencing progeny skew. We used a multiple-merger coalescent model of progeny skew (Irwin et al. 2016) which required two parameters: *Ξ* corresponding to the probability of a virion having multiple reproduction events in a single tick, and the size of this reproductive burst. Values of *Ξ* represent the eclipse time of a virion (10 hours, *i.e.*, a value of *Ξ* = 0.1) as the max rate (Bar-On et al. 2020) or the product of this eclipse time and the number of virions present in a cell that is actively budding virions (*i.e.*, 100000 virions multiplied by 10 hours, or a value of *Ξ* = 0.000001) as the minimum rate (Sender et al. 2021). The geometric mean of the minimum and maximum value (0.001) was used as the midpoint value. We selected the highest level of burst size by determining the largest possible burst that could still be completed with the maximum levels of *Ξ* and carrying capacity and the requested computing resources; we selected lower levels as proportions of that maximum value. Finally, we parameterized the DFE as the ratio of strongly deleterious relative to neutral mutations.

Simulations used a common, burn-in source population which was created using a full-model simulation run for 25,000 ticks at the highest value of carrying capacity and the mid-level value for all other parameters. We replicated each parameter combination 5 times. Therefore, we ran a total of 12,096 model-parameter combinations (represented visually in Figure 1), resulting in 60,480 replicates. To simulate variation in sequencing depth of empirical studies, we independently sampled 100 and 1000 genomes and stored them as .ms files. A total of 26 replicates failed to complete due to requiring more than the allotted computational resources and were excluded from further analysis (Supplemental Table 2).

### Simulation Assessment

Using the scikit-allel package (Miles et al. 2021) and custom python script (Rossum and Drake 2009), we calculated a set of summary statistics for each .ms file. The primary statistic we used to identify models resulting in levels of variation similar to that observed in empirical patient samples was the total number of SNPs (often referred to as intra-host single nucleotide variations or iSNVs in the SARS-CoV-2 literature). We compared the simulated data to multiple recent studies that have sought to quantify the amount of intra-host variation in SARS-CoV-2 (Lythgoe et al. 2021, Valesano et al. 2021, Wang et al. 2021, Bendall et al. 2023, Gu et al. 2023). To make this comparison, we applied filtering criteria to the simulated data reflecting that implemented in the empirical data (*i.e.,* removing variants segregating below 2% frequency). From this existing literature, the number of SNPs found within an intra-host population of SARS-CoV-2 clearing this frequency threshold tends to be 5 or fewer (Table 4). Additionally, we required each parameter combination to clear this threshold for both the 100 and 1000 genome sampling schemes. We performed model filtering, and figure creation was performed using a custom R script (R Core Team 2022) and the packages ggplot2 (Wickham 2016) and gridExtra (Auguie 2017). All eidos, bash, python, and R scripts have been deposited on github (https://github.com/jwterbot2/SARS-CoV-2_InitialBaselineModel)

## Supplementary Material

Supplementary tables and figures are available in a linked file for the purposes of review.

## Acknowledgements

Research reported in this manuscript was supported by the National Institute of General Medical Sciences of the National Institutes of Health (NIH) under Award Number P30GM140963 (to JMG and BSC). NIH awards R35GM124701 (BSC) and R35GM139383 (JDJ) also supported this work. The funders had no role in study design, data collection and analysis, decision to publish, or preparation of the manuscript. We are grateful to Parul Johri and Vivak Soni for their advice regarding simulation design. We would also like to thank the computing resources used during this study: the common burn-in was run on the Agave computing cluster at ASU and simulation replicates were run using the Open Science Pool resources (OSG 2006, Pordes et al. 2007, Sfiligoi et al. 2009).

## Data Availability

Scripts and code used in this study along with a copy of data used to generate figures are available online via github (https://github.com/jwterbot2/SARS-CoV-2_InitialBaselineModel).

---

**Supplemental Table 1.**
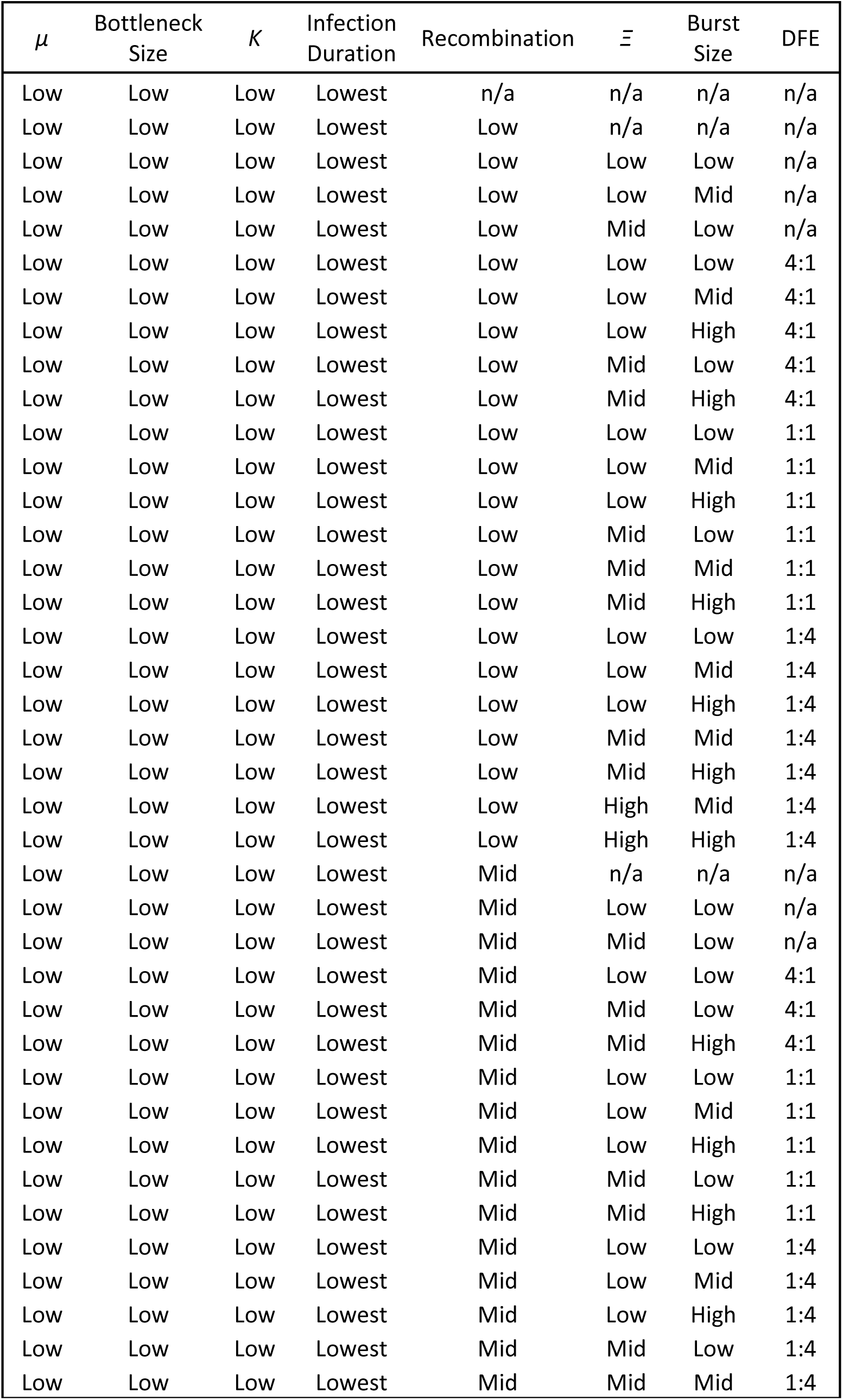
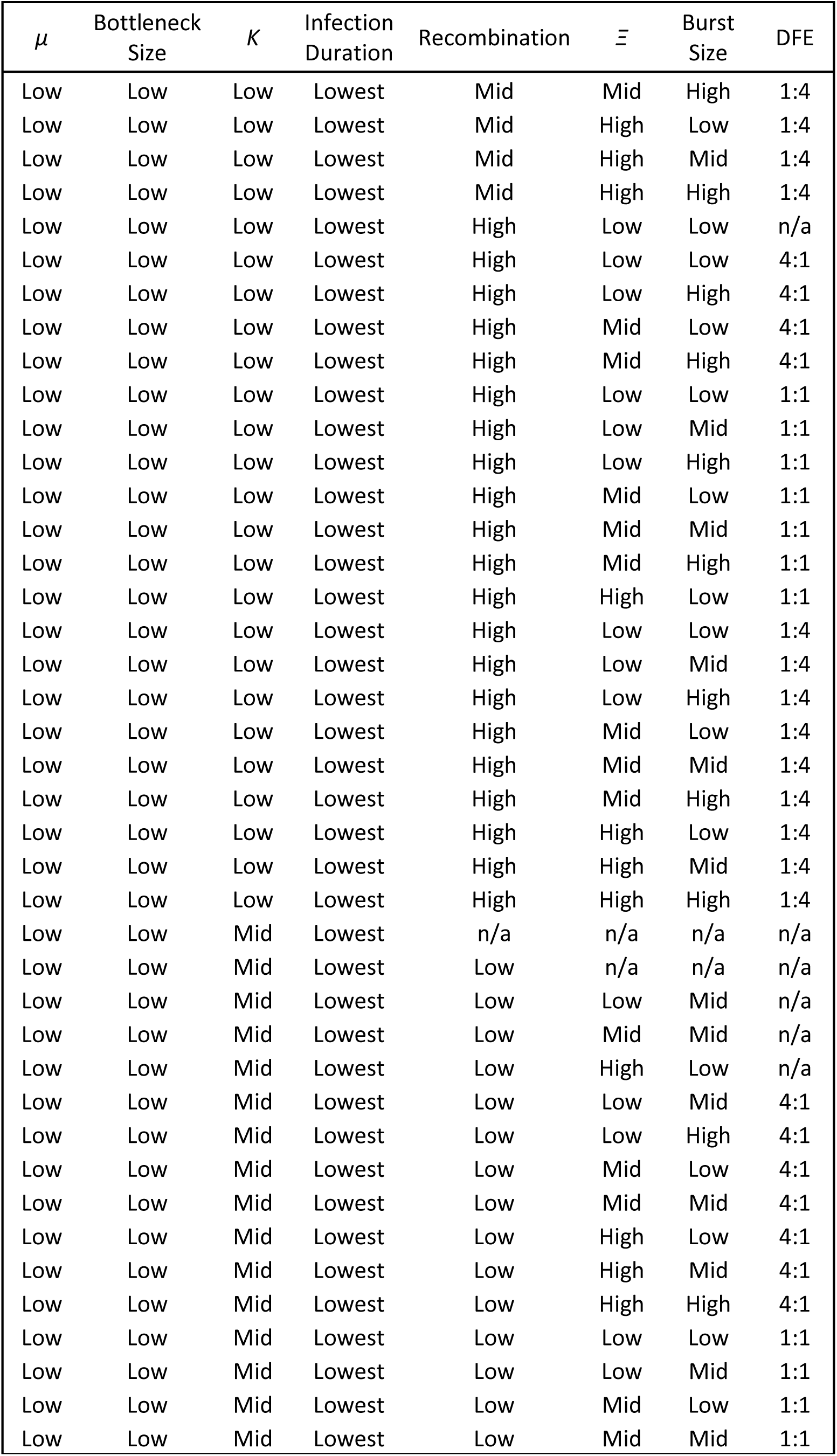
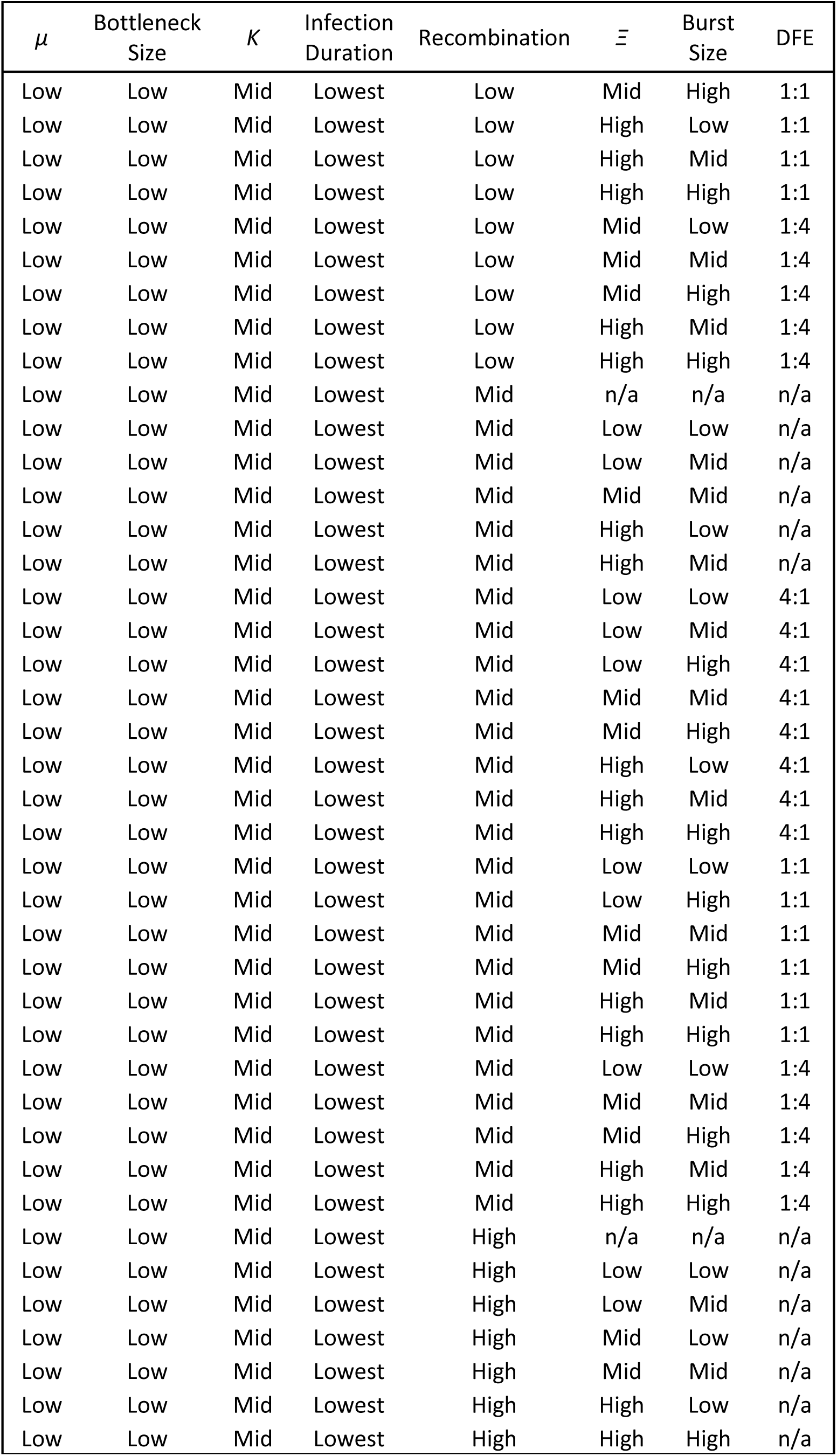
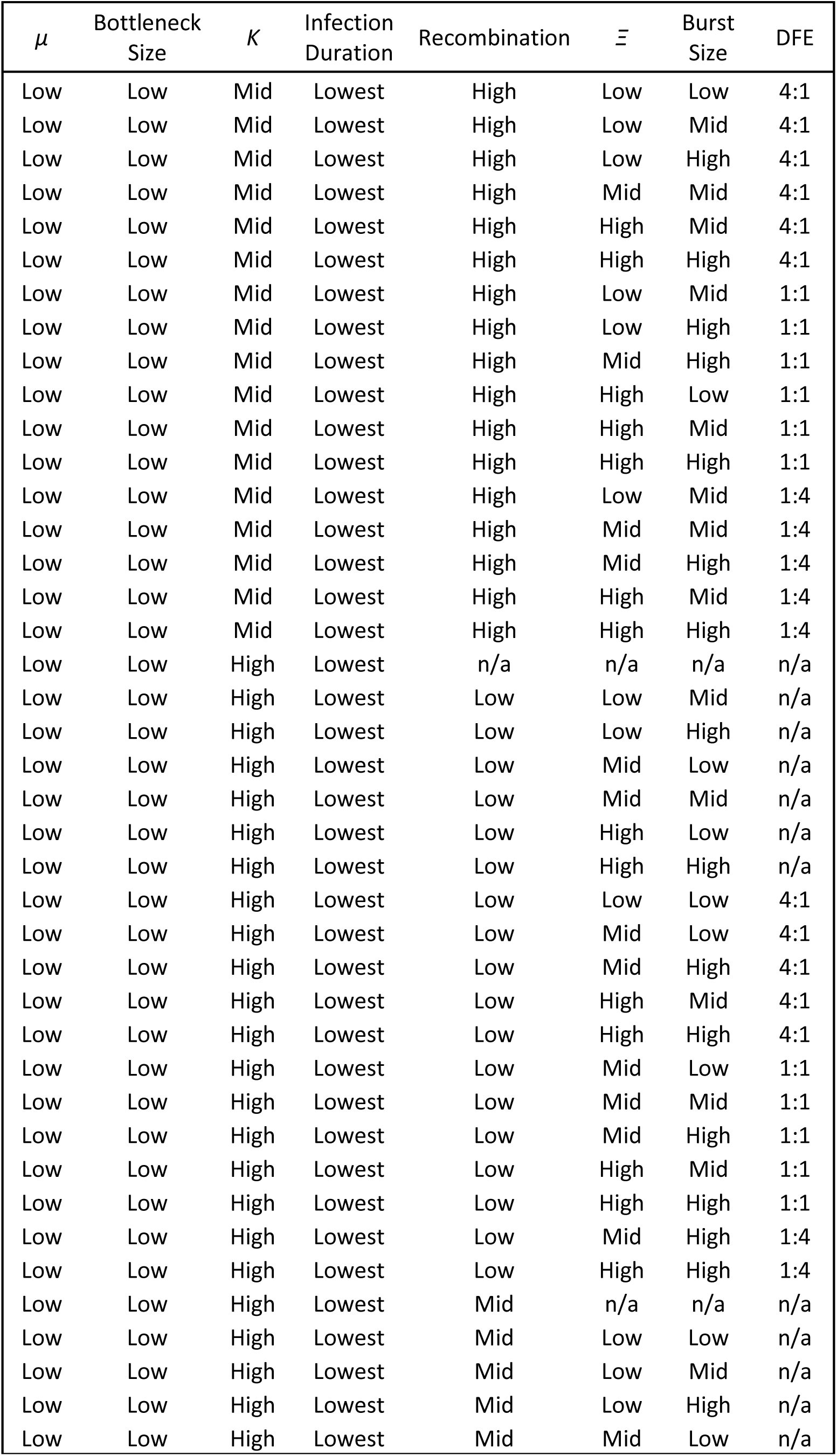
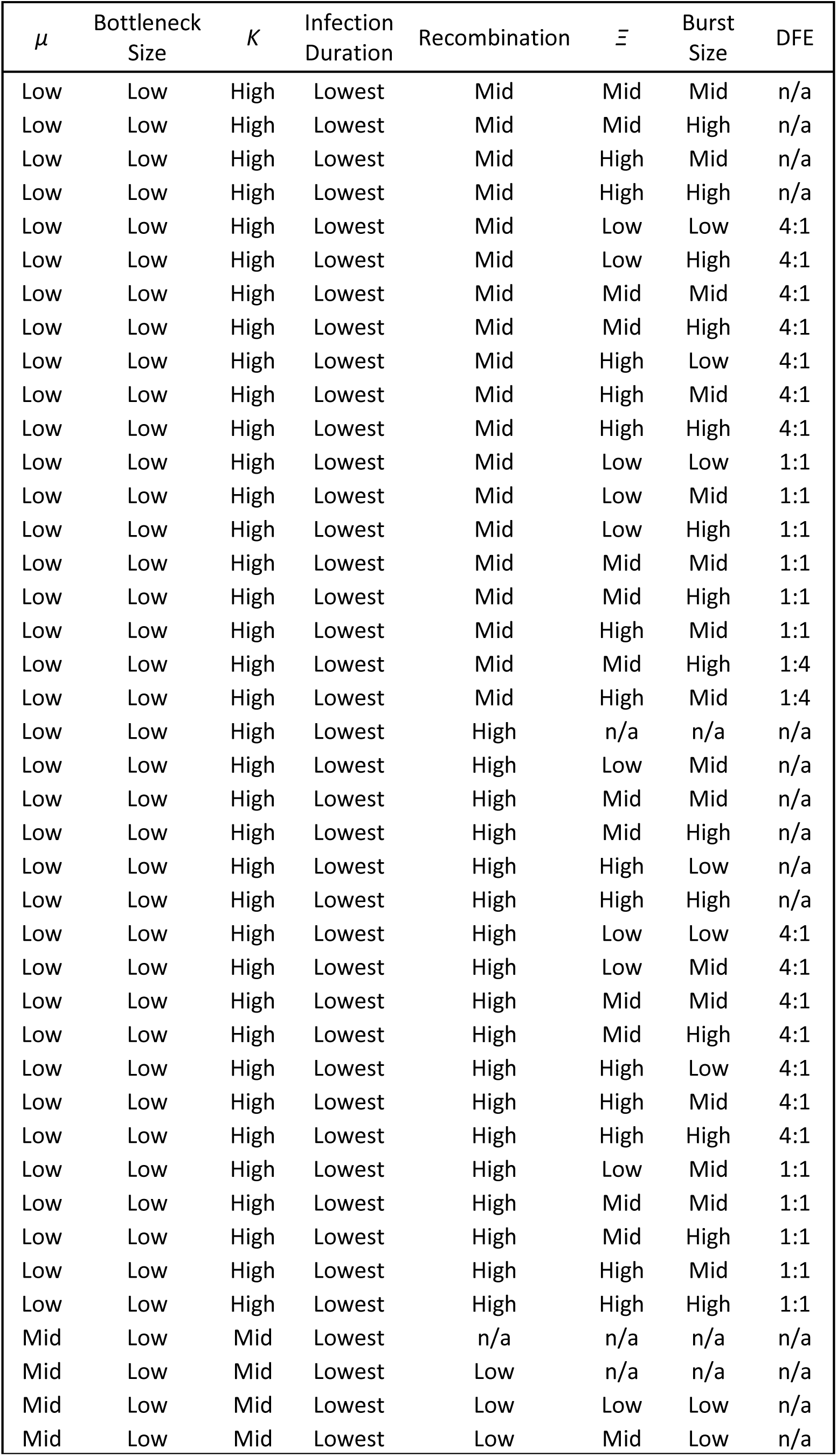
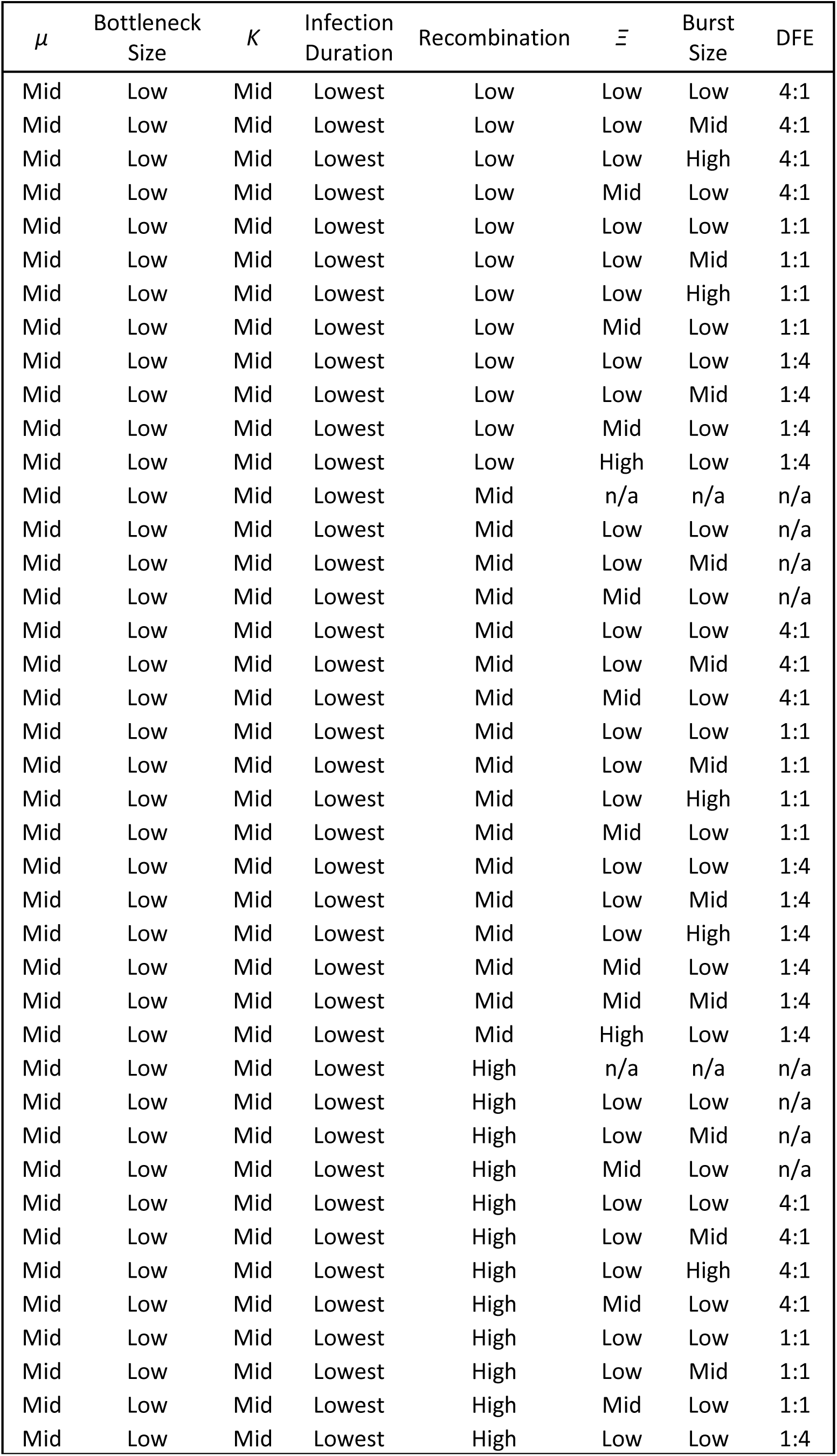
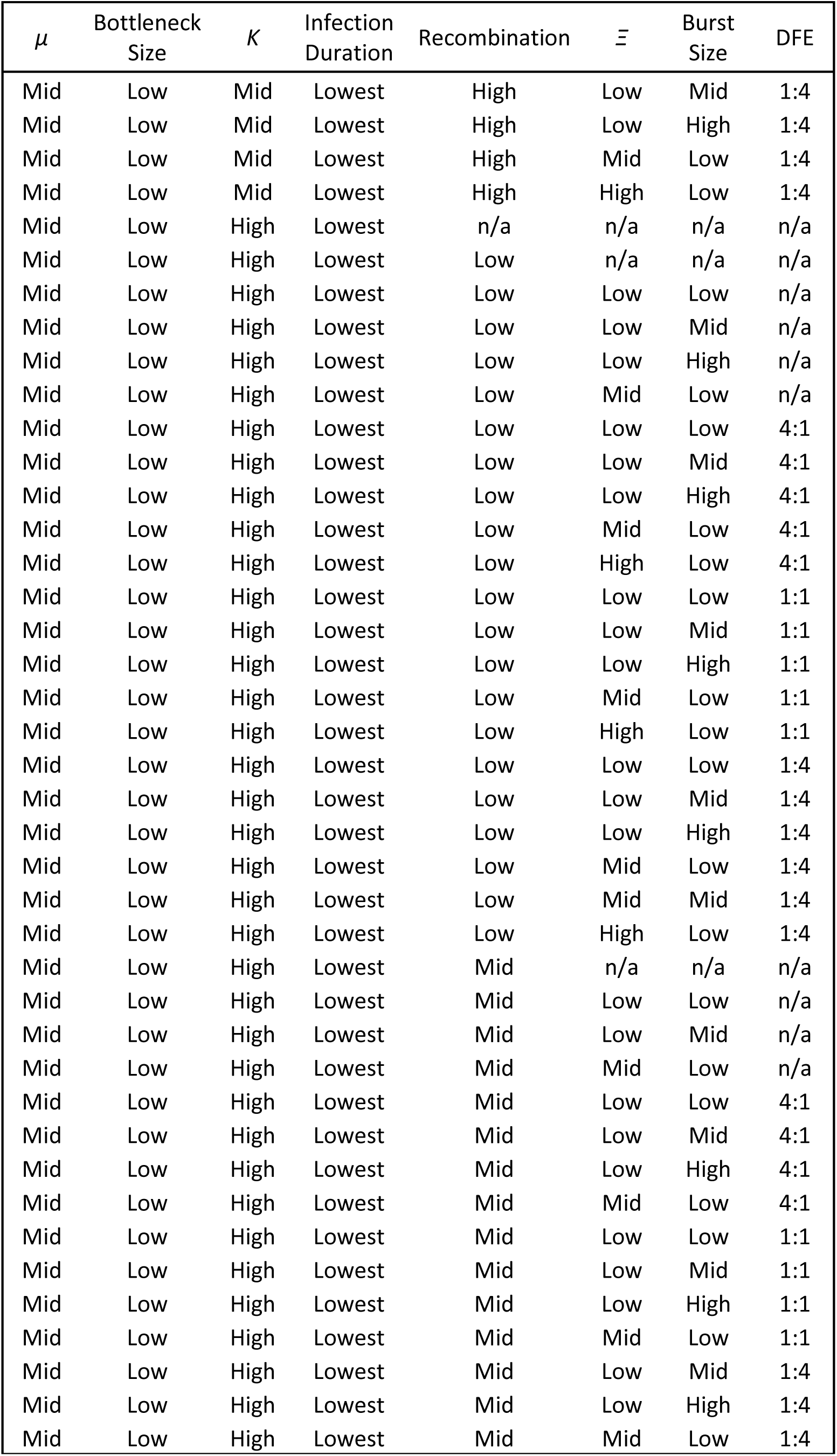
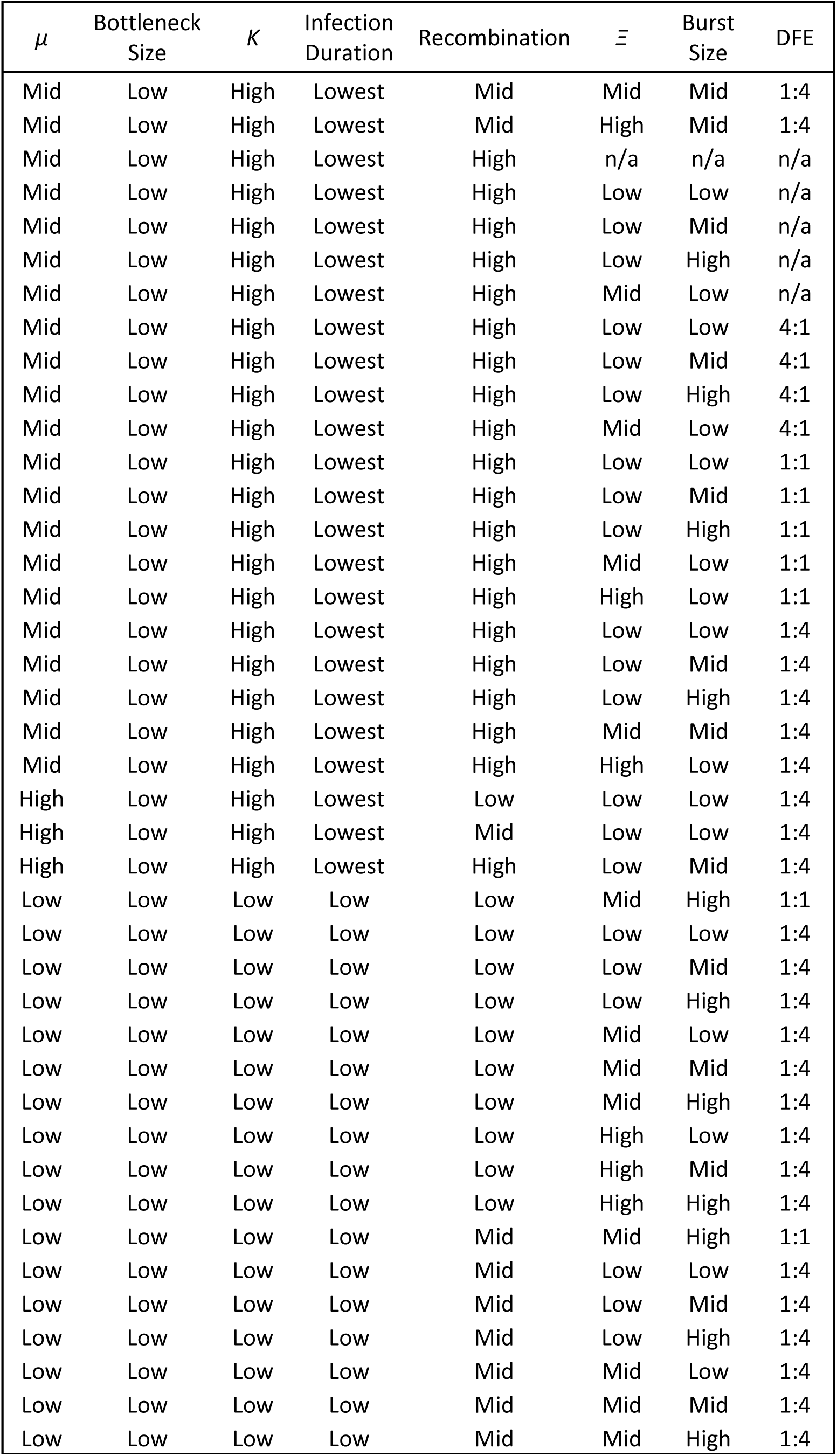
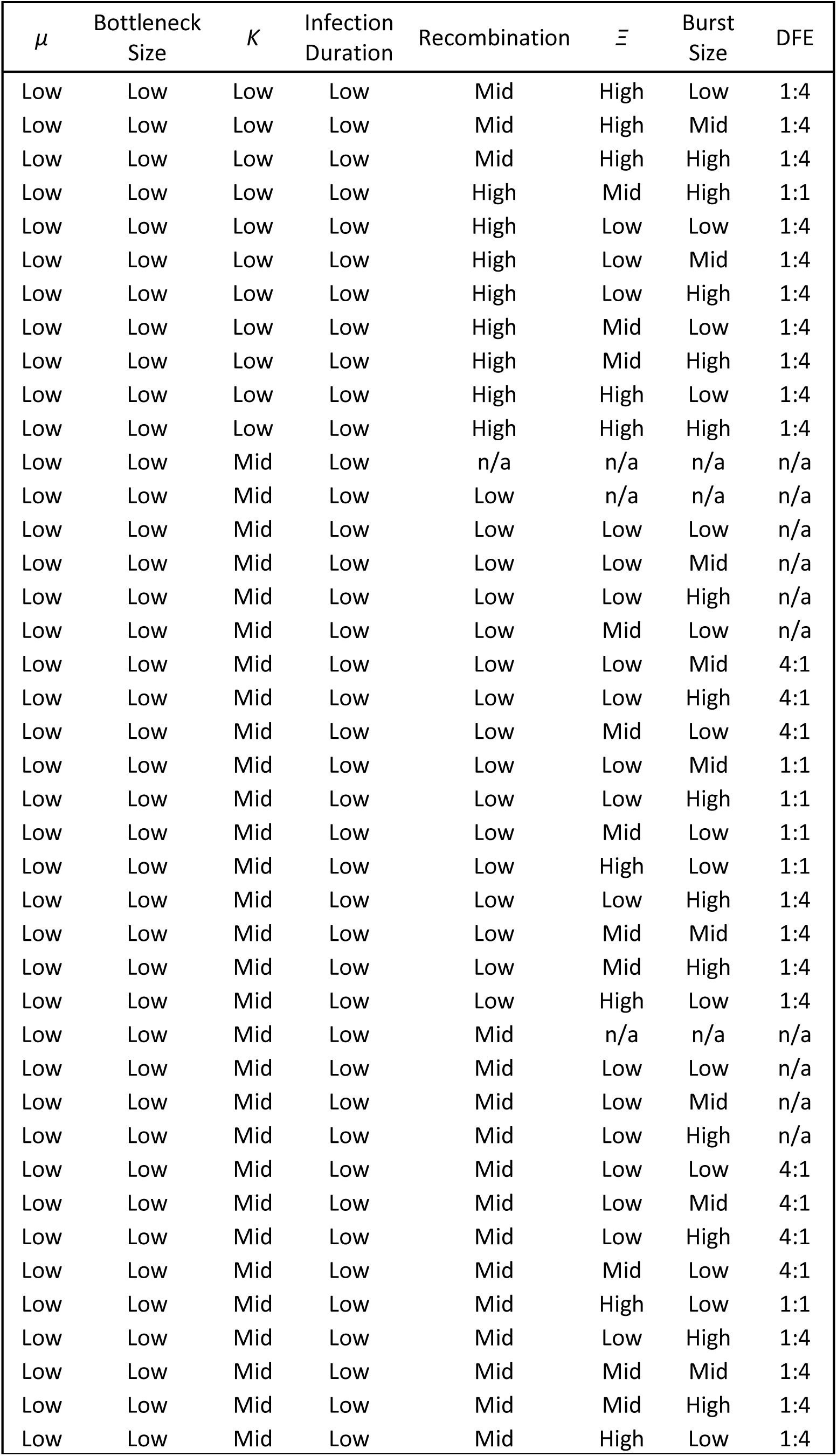
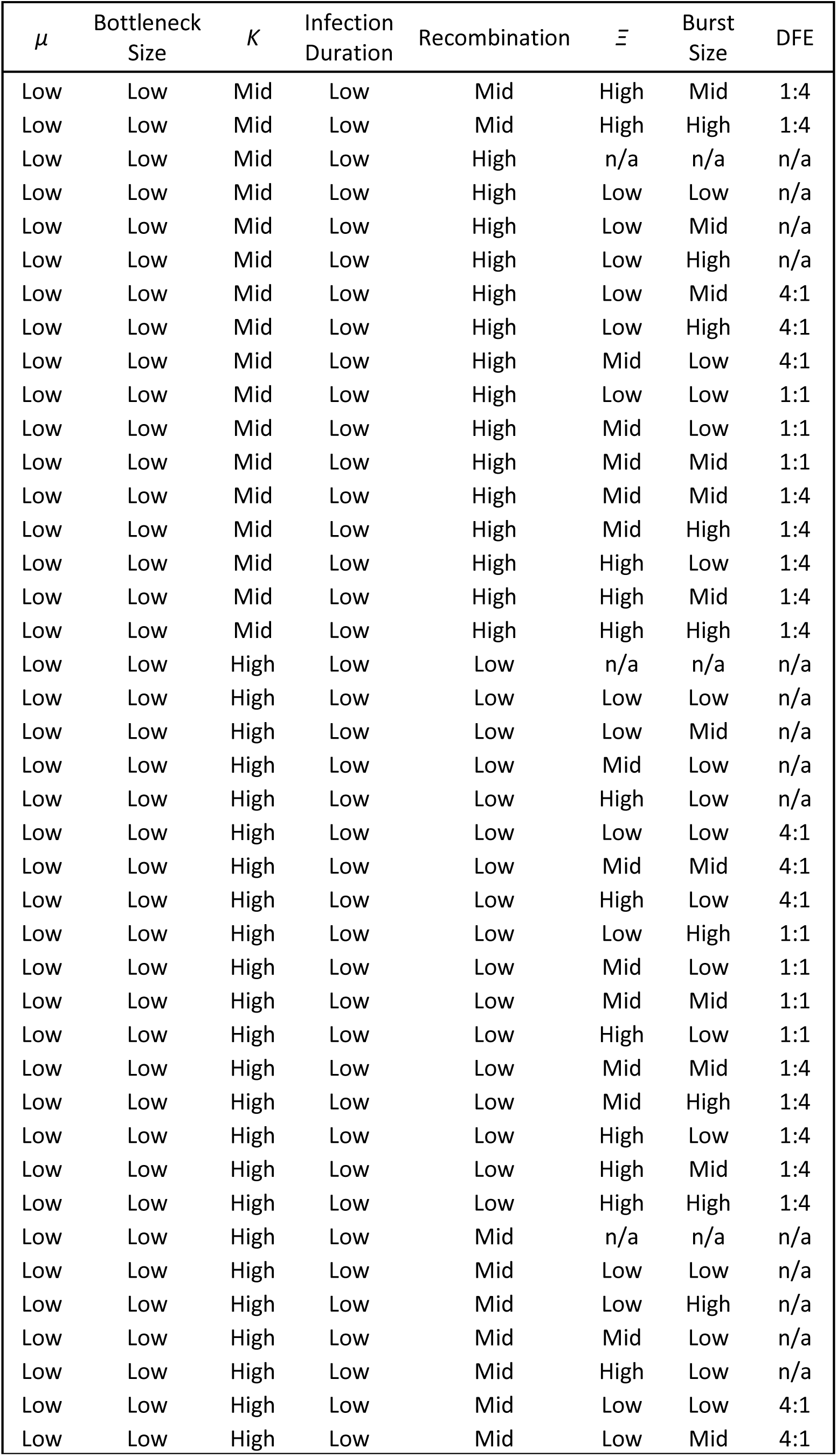
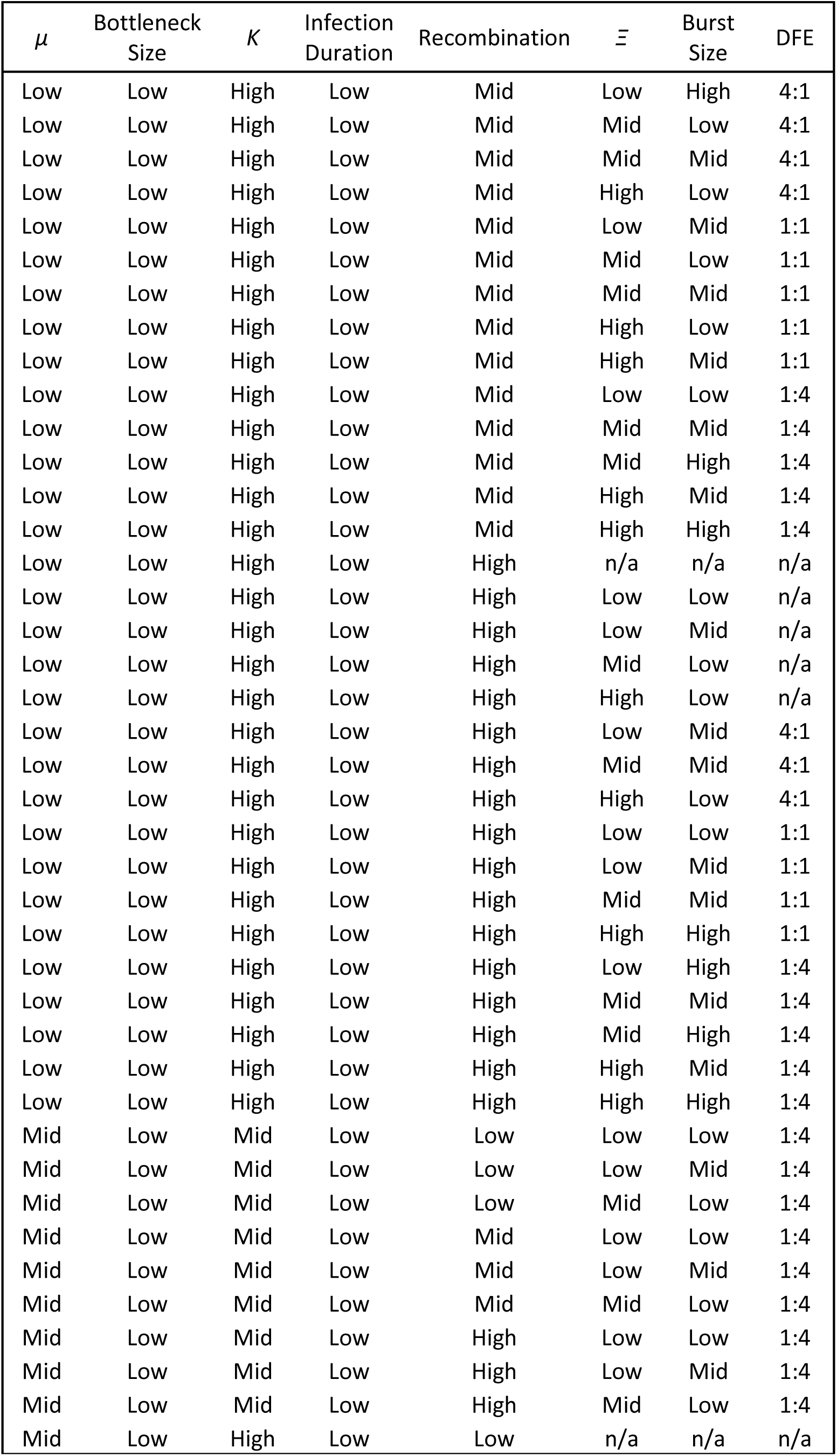
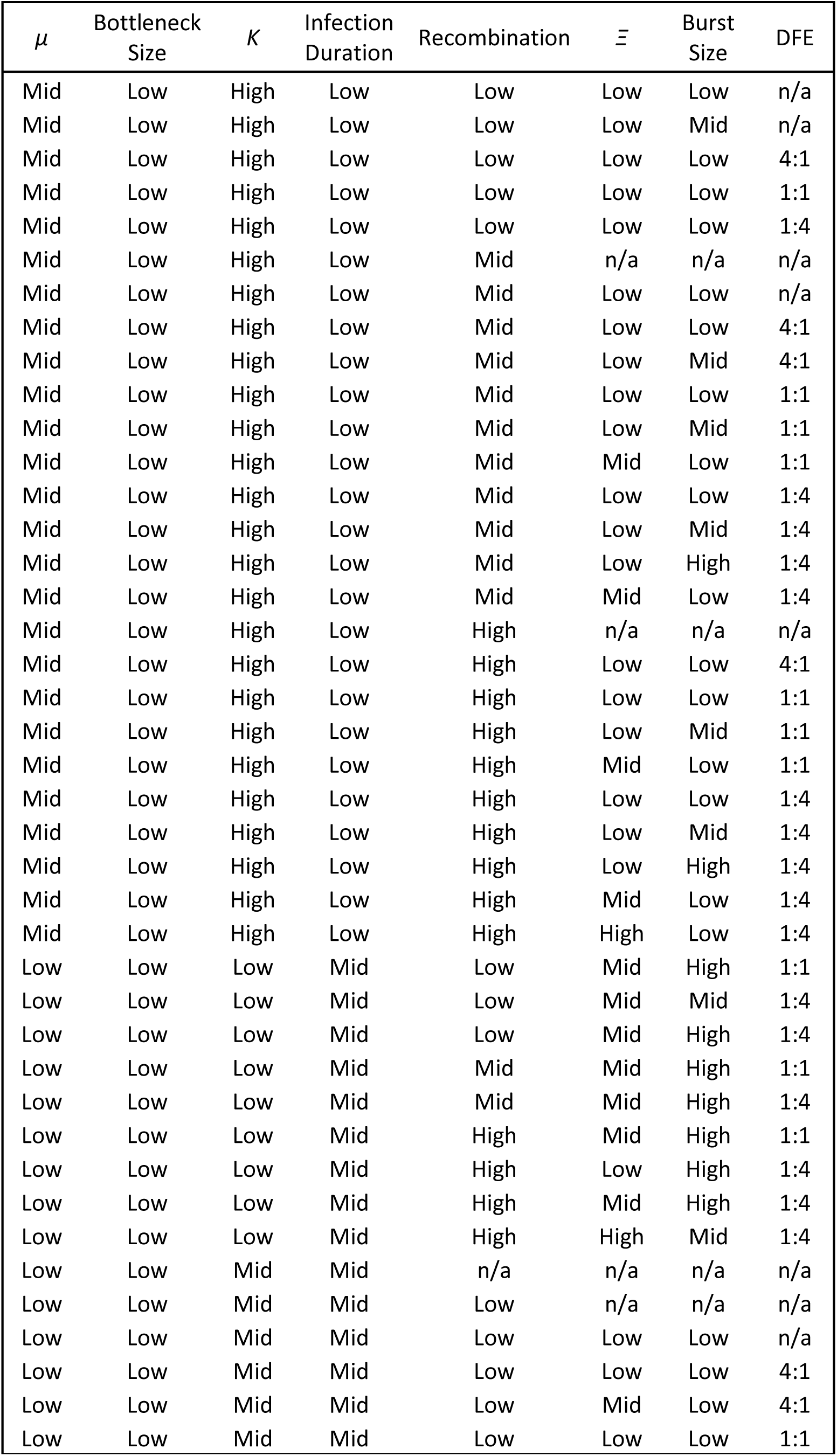
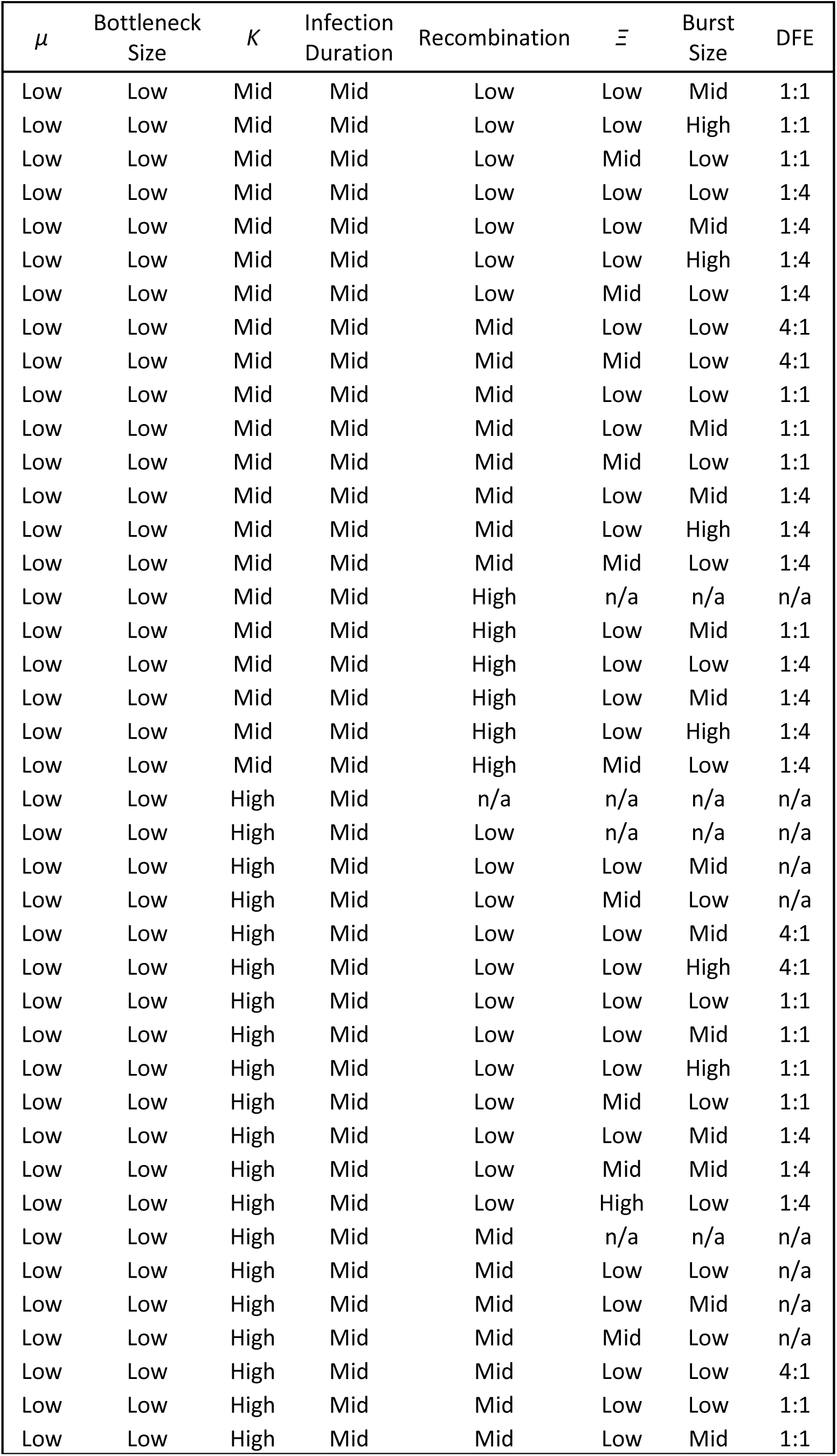
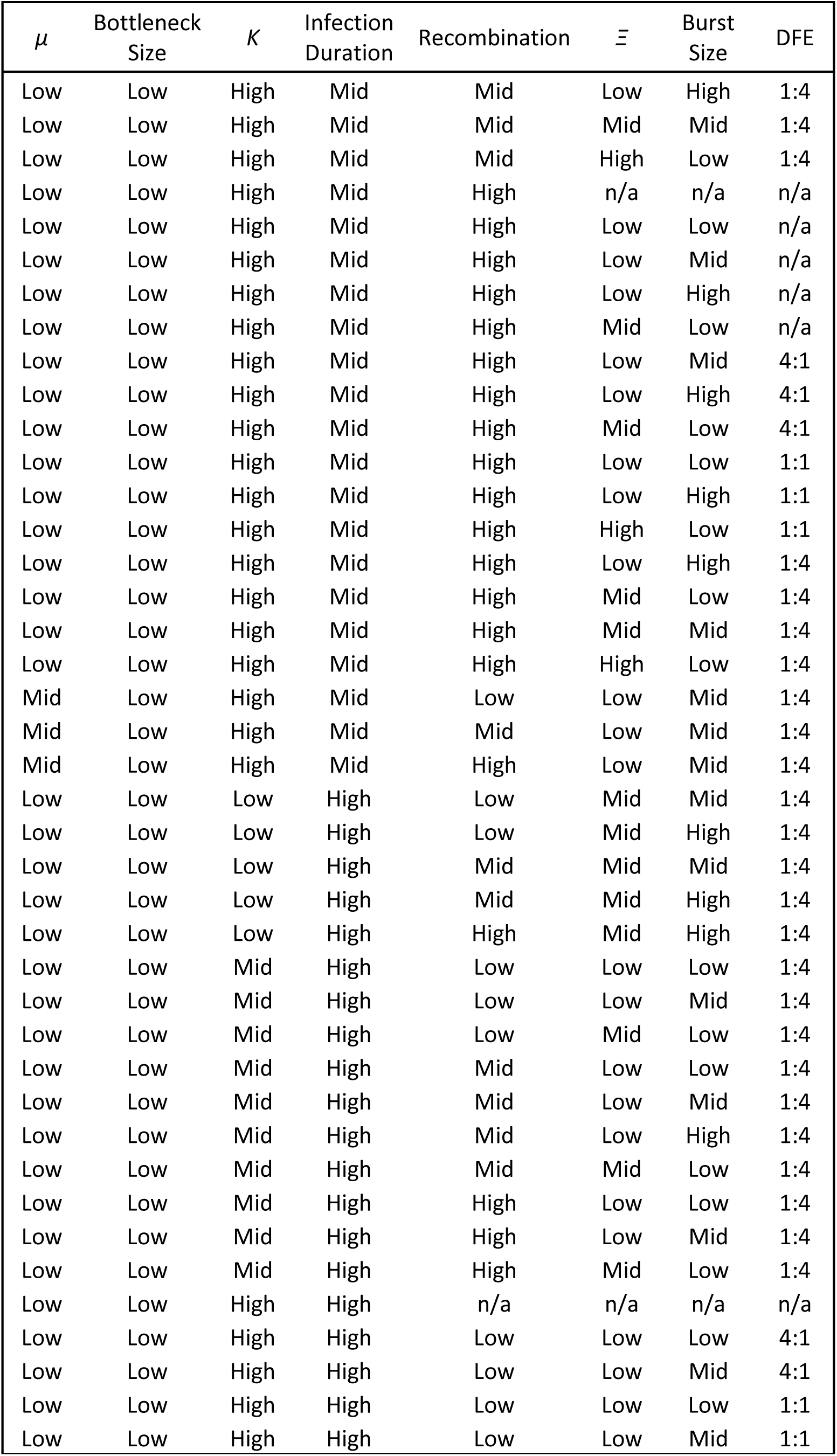
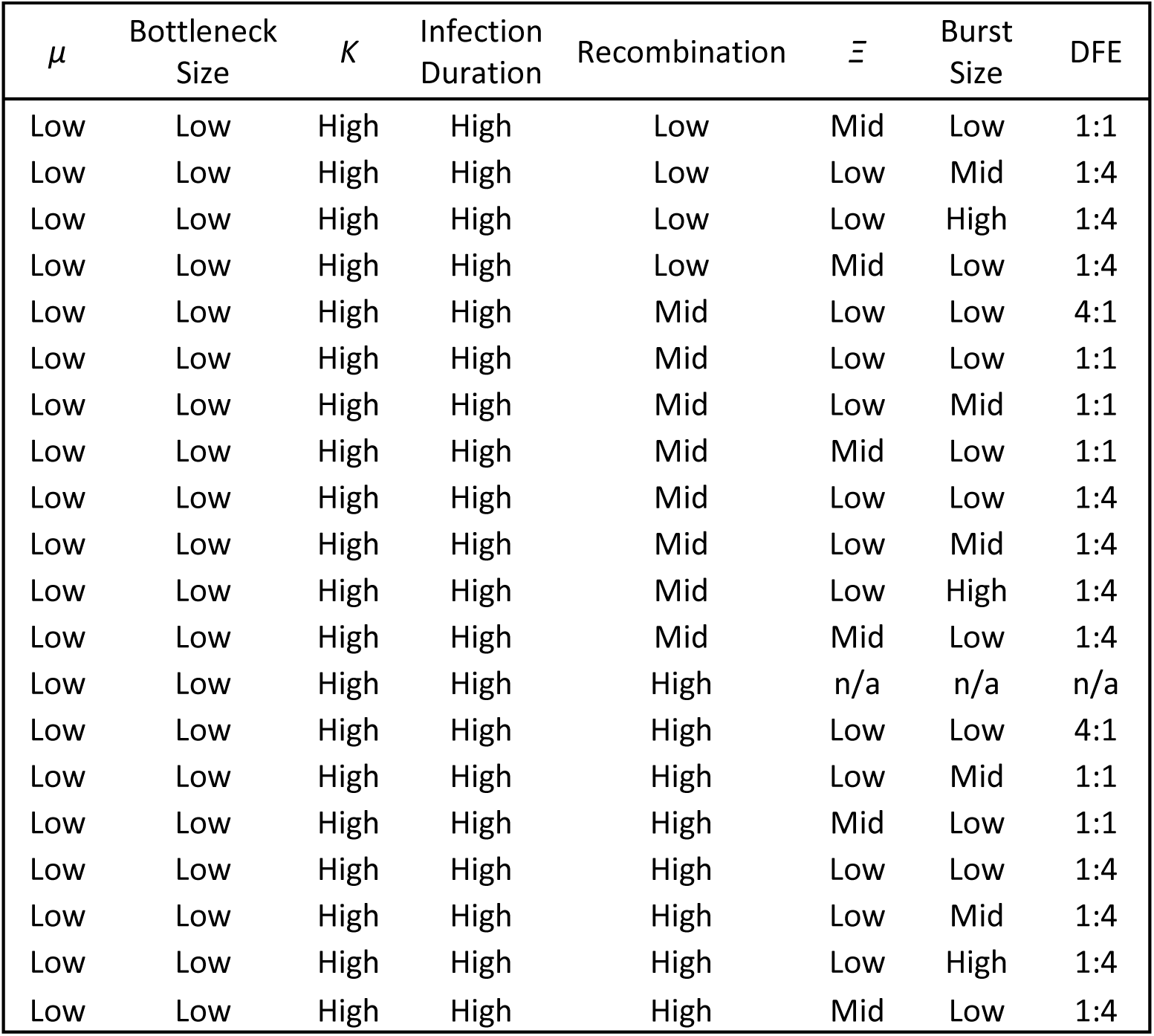
List of parameter levels for all plausible models.

| $\mu$ | Bottleneck Size | $K$ | Infection Duration | Recombination | $\Xi$ | Burst Size | DFE |
| --- | --- | --- | --- | --- | --- | --- | --- |
| Low | Low | Low | Lowest | n/a | n/a | n/a | n/a |
| Low | Low | Low | Lowest | Low | n/a | n/a | n/a |
| Low | Low | Low | Lowest | Low | Low | Low | n/a |
| Low | Low | Low | Lowest | Low | Low | Mid | n/a |
| Low | Low | Low | Lowest | Low | Mid | Low | n/a |
| Low | Low | Low | Lowest | Low | Low | Low | 4:1 |
| Low | Low | Low | Lowest | Low | Low | Mid | 4:1 |
| Low | Low | Low | Lowest | Low | Low | High | 4:1 |
| Low | Low | Low | Lowest | Low | Mid | Low | 4:1 |
| Low | Low | Low | Lowest | Low | Mid | High | 4:1 |
| Low | Low | Low | Lowest | Low | Low | Low | 1:1 |
| Low | Low | Low | Lowest | Low | Low | Mid | 1:1 |
| Low | Low | Low | Lowest | Low | Low | High | 1:1 |
| Low | Low | Low | Lowest | Low | Mid | Low | 1:1 |
| Low | Low | Low | Lowest | Low | Mid | Mid | 1:1 |
| Low | Low | Low | Lowest | Low | Mid | High | 1:1 |
| Low | Low | Low | Lowest | Low | Low | Low | 1:4 |
| Low | Low | Low | Lowest | Low | Low | Mid | 1:4 |
| Low | Low | Low | Lowest | Low | Low | High | 1:4 |
| Low | Low | Low | Lowest | Low | Mid | Mid | 1:4 |
| Low | Low | Low | Lowest | Low | Mid | High | 1:4 |
| Low | Low | Low | Lowest | Low | High | Mid | 1:4 |
| Low | Low | Low | Lowest | Low | High | High | 1:4 |
| Low | Low | Low | Lowest | Mid | n/a | n/a | n/a |
| Low | Low | Low | Lowest | Mid | Low | Low | n/a |
| Low | Low | Low | Lowest | Mid | Mid | Low | n/a |
| Low | Low | Low | Lowest | Mid | Low | Low | 4:1 |
| Low | Low | Low | Lowest | Mid | Mid | Low | 4:1 |
| Low | Low | Low | Lowest | Mid | Mid | High | 4:1 |
| Low | Low | Low | Lowest | Mid | Low | Low | 1:1 |
| Low | Low | Low | Lowest | Mid | Low | Mid | 1:1 |
| Low | Low | Low | Lowest | Mid | Low | High | 1:1 |
| Low | Low | Low | Lowest | Mid | Mid | Low | 1:1 |
| Low | Low | Low | Lowest | Mid | Mid | High | 1:1 |
| Low | Low | Low | Lowest | Mid | Low | Low | 1:4 |
| Low | Low | Low | Lowest | Mid | Low | Mid | 1:4 |
| Low | Low | Low | Lowest | Mid | Low | High | 1:4 |
| Low | Low | Low | Lowest | Mid | Mid | Low | 1:4 |
| Low | Low | Low | Lowest | Mid | Mid | Mid | 1:4 |
| Low | Low | Low | Lowest | Mid | Mid | High | 1:4 |
| Low | Low | Low | Lowest | Mid | High | Low | 1:4 |
| Low | Low | Low | Lowest | Mid | High | Mid | 1:4 |
| Low | Low | Low | Lowest | Mid | High | High | 1:4 |
| Low | Low | Low | Lowest | High | Low | Low | n/a |
| Low | Low | Low | Lowest | High | Low | Low | 4:1 |
| Low | Low | Low | Lowest | High | Low | High | 4:1 |
| Low | Low | Low | Lowest | High | Mid | Low | 4:1 |
| Low | Low | Low | Lowest | High | Mid | High | 4:1 |
| Low | Low | Low | Lowest | High | Low | Low | 1:1 |
| Low | Low | Low | Lowest | High | Low | Mid | 1:1 |
| Low | Low | Low | Lowest | High | Low | High | 1:1 |
| Low | Low | Low | Lowest | High | Mid | Low | 1:1 |
| Low | Low | Low | Lowest | High | Mid | Mid | 1:1 |
| Low | Low | Low | Lowest | High | Mid | High | 1:1 |
| Low | Low | Low | Lowest | High | High | Low | 1:1 |
| Low | Low | Low | Lowest | High | Low | Low | 1:4 |
| Low | Low | Low | Lowest | High | Low | Mid | 1:4 |
| Low | Low | Low | Lowest | High | Low | High | 1:4 |
| Low | Low | Low | Lowest | High | Mid | Low | 1:4 |
| Low | Low | Low | Lowest | High | Mid | Mid | 1:4 |
| Low | Low | Low | Lowest | High | Mid | High | 1:4 |
| Low | Low | Low | Lowest | High | High | Low | 1:4 |
| Low | Low | Low | Lowest | High | High | Mid | 1:4 |
| Low | Low | Low | Lowest | High | High | High | 1:4 |
| Low | Low | Mid | Lowest | n/a | n/a | n/a | n/a |
| Low | Low | Mid | Lowest | Low | n/a | n/a | n/a |
| Low | Low | Mid | Lowest | Low | Low | Mid | n/a |
| Low | Low | Mid | Lowest | Low | Mid | Mid | n/a |
| Low | Low | Mid | Lowest | Low | High | Low | n/a |
| Low | Low | Mid | Lowest | Low | Low | Mid | 4:1 |
| Low | Low | Mid | Lowest | Low | Low | High | 4:1 |
| Low | Low | Mid | Lowest | Low | Mid | Low | 4:1 |
| Low | Low | Mid | Lowest | Low | Mid | Mid | 4:1 |
| Low | Low | Mid | Lowest | Low | High | Low | 4:1 |
| Low | Low | Mid | Lowest | Low | High | Mid | 4:1 |
| Low | Low | Mid | Lowest | Low | High | High | 4:1 |
| Low | Low | Mid | Lowest | Low | Low | Low | 1:1 |
| Low | Low | Mid | Lowest | Low | Low | Mid | 1:1 |
| Low | Low | Mid | Lowest | Low | Mid | Low | 1:1 |
| Low | Low | Mid | Lowest | Low | Mid | Mid | 1:1 |
| Low | Low | Mid | Lowest | Low | Mid | High | 1:1 |
| Low | Low | Mid | Lowest | Low | High | Low | 1:1 |
| Low | Low | Mid | Lowest | Low | High | Mid | 1:1 |
| Low | Low | Mid | Lowest | Low | High | High | 1:1 |
| Low | Low | Mid | Lowest | Low | Mid | Low | 1:4 |
| Low | Low | Mid | Lowest | Low | Mid | Mid | 1:4 |
| Low | Low | Mid | Lowest | Low | Mid | High | 1:4 |
| Low | Low | Mid | Lowest | Low | High | Mid | 1:4 |
| Low | Low | Mid | Lowest | Low | High | High | 1:4 |
| Low | Low | Mid | Lowest | Mid | n/a | n/a | n/a |
| Low | Low | Mid | Lowest | Mid | Low | Low | n/a |
| Low | Low | Mid | Lowest | Mid | Low | Mid | n/a |
| Low | Low | Mid | Lowest | Mid | Mid | Mid | n/a |
| Low | Low | Mid | Lowest | Mid | High | Low | n/a |
| Low | Low | Mid | Lowest | Mid | High | Mid | n/a |
| Low | Low | Mid | Lowest | Mid | Low | Low | 4:1 |
| Low | Low | Mid | Lowest | Mid | Low | Mid | 4:1 |
| Low | Low | Mid | Lowest | Mid | Low | High | 4:1 |
| Low | Low | Mid | Lowest | Mid | Mid | Mid | 4:1 |
| Low | Low | Mid | Lowest | Mid | Mid | High | 4:1 |
| Low | Low | Mid | Lowest | Mid | High | Low | 4:1 |
| Low | Low | Mid | Lowest | Mid | High | Mid | 4:1 |
| Low | Low | Mid | Lowest | Mid | High | High | 4:1 |
| Low | Low | Mid | Lowest | Mid | Low | Low | 1:1 |
| Low | Low | Mid | Lowest | Mid | Low | High | 1:1 |
| Low | Low | Mid | Lowest | Mid | Mid | Mid | 1:1 |
| Low | Low | Mid | Lowest | Mid | Mid | High | 1:1 |
| Low | Low | Mid | Lowest | Mid | High | Mid | 1:1 |
| Low | Low | Mid | Lowest | Mid | High | High | 1:1 |
| Low | Low | Mid | Lowest | Mid | Low | Low | 1:4 |
| Low | Low | Mid | Lowest | Mid | Mid | Mid | 1:4 |
| Low | Low | Mid | Lowest | Mid | Mid | High | 1:4 |
| Low | Low | Mid | Lowest | Mid | High | Mid | 1:4 |
| Low | Low | Mid | Lowest | Mid | High | High | 1:4 |
| Low | Low | Mid | Lowest | High | n/a | n/a | n/a |
| Low | Low | Mid | Lowest | High | Low | Low | n/a |
| Low | Low | Mid | Lowest | High | Low | Mid | n/a |
| Low | Low | Mid | Lowest | High | Mid | Low | n/a |
| Low | Low | Mid | Lowest | High | Mid | Mid | n/a |
| Low | Low | Mid | Lowest | High | High | Low | n/a |
| Low | Low | Mid | Lowest | High | High | High | n/a |

| $\mu$ | Bottleneck Size | $K$ | Infection Duration | Recombination | $\Xi$ | Burst Size | DfE |
| --- | --- | --- | --- | --- | --- | --- | --- |
| Low | Low | Mid | Lowest | High | Low | Low | 4:1 |
| Low | Low | Mid | Lowest | High | Low | Mid | 4:1 |
| Low | Low | Mid | Lowest | High | Low | High | 4:1 |
| Low | Low | Mid | Lowest | High | Mid | Mid | 4:1 |
| Low | Low | Mid | Lowest | High | High | Mid | 4:1 |
| Low | Low | Mid | Lowest | High | High | High | 4:1 |
| Low | Low | Mid | Lowest | High | Low | Mid | 1:1 |
| Low | Low | Mid | Lowest | High | Low | High | 1:1 |
| Low | Low | Mid | Lowest | High | Mid | High | 1:1 |
| Low | Low | Mid | Lowest | High | High | Low | 1:1 |
| Low | Low | Mid | Lowest | High | High | Mid | 1:1 |
| Low | Low | Mid | Lowest | High | High | High | 1:1 |
| Low | Low | Mid | Lowest | High | Low | Mid | 1:4 |
| Low | Low | Mid | Lowest | High | Mid | Mid | 1:4 |
| Low | Low | Mid | Lowest | High | Mid | High | 1:4 |
| Low | Low | Mid | Lowest | High | High | Mid | 1:4 |
| Low | Low | Mid | Lowest | High | High | High | 1:4 |
| Low | Low | High | Lowest | n/a | n/a | n/a | n/a |
| Low | Low | High | Lowest | Low | Low | Mid | n/a |
| Low | Low | High | Lowest | Low | Low | High | n/a |
| Low | Low | High | Lowest | Low | Mid | Low | n/a |
| Low | Low | High | Lowest | Low | Mid | Mid | n/a |
| Low | Low | High | Lowest | Low | High | Low | n/a |
| Low | Low | High | Lowest | Low | High | High | n/a |
| Low | Low | High | Lowest | Low | Low | Low | 4:1 |
| Low | Low | High | Lowest | Low | Mid | Low | 4:1 |
| Low | Low | High | Lowest | Low | Mid | High | 4:1 |
| Low | Low | High | Lowest | Low | High | Mid | 4:1 |
| Low | Low | High | Lowest | Low | High | High | 4:1 |
| Low | Low | High | Lowest | Low | Mid | Low | 1:1 |
| Low | Low | High | Lowest | Low | Mid | Mid | 1:1 |
| Low | Low | High | Lowest | Low | Mid | High | 1:1 |
| Low | Low | High | Lowest | Low | High | Mid | 1:1 |
| Low | Low | High | Lowest | Low | High | High | 1:1 |
| Low | Low | High | Lowest | Low | Mid | High | 1:4 |
| Low | Low | High | Lowest | Low | High | High | 1:4 |
| Low | Low | High | Lowest | Mid | n/a | n/a | n/a |
| Low | Low | High | Lowest | Mid | Low | Low | n/a |
| Low | Low | High | Lowest | Mid | Low | Mid | n/a |
| Low | Low | High | Lowest | Mid | Low | High | n/a |
| Low | Low | High | Lowest | Mid | Mid | Low | n/a |
| Low | Low | High | Lowest | Mid | Mid | Mid | n/a |
| Low | Low | High | Lowest | Mid | Mid | High | n/a |
| Low | Low | High | Lowest | Mid | High | Mid | n/a |
| Low | Low | High | Lowest | Mid | High | High | n/a |
| Low | Low | High | Lowest | Mid | Low | Low | 4:1 |
| Low | Low | High | Lowest | Mid | Low | High | 4:1 |
| Low | Low | High | Lowest | Mid | Mid | Mid | 4:1 |
| Low | Low | High | Lowest | Mid | Mid | High | 4:1 |
| Low | Low | High | Lowest | Mid | High | Low | 4:1 |
| Low | Low | High | Lowest | Mid | High | Mid | 4:1 |
| Low | Low | High | Lowest | Mid | High | High | 4:1 |
| Low | Low | High | Lowest | Mid | Low | Low | 1:1 |
| Low | Low | High | Lowest | Mid | Low | Mid | 1:1 |
| Low | Low | High | Lowest | Mid | Low | High | 1:1 |
| Low | Low | High | Lowest | Mid | Mid | Mid | 1:1 |
| Low | Low | High | Lowest | Mid | Mid | High | 1:1 |
| Low | Low | High | Lowest | Mid | High | Mid | 1:1 |
| Low | Low | High | Lowest | Mid | Mid | High | 1:4 |
| Low | Low | High | Lowest | Mid | High | Mid | 1:4 |
| Low | Low | High | Lowest | High | n/a | n/a | n/a |
| Low | Low | High | Lowest | High | Low | Mid | n/a |
| Low | Low | High | Lowest | High | Mid | Mid | n/a |
| Low | Low | High | Lowest | High | Mid | High | n/a |
| Low | Low | High | Lowest | High | High | Low | n/a |
| Low | Low | High | Lowest | High | High | High | n/a |
| Low | Low | High | Lowest | High | Low | Low | 4:1 |
| Low | Low | High | Lowest | High | Low | Mid | 4:1 |
| Low | Low | High | Lowest | High | Mid | Mid | 4:1 |
| Low | Low | High | Lowest | High | Mid | High | 4:1 |
| Low | Low | High | Lowest | High | High | Low | 4:1 |
| Low | Low | High | Lowest | High | High | Mid | 4:1 |
| Low | Low | High | Lowest | High | High | High | 4:1 |
| Low | Low | High | Lowest | High | Low | Mid | 1:1 |
| Low | Low | High | Lowest | High | Mid | Mid | 1:1 |
| Low | Low | High | Lowest | High | Mid | High | 1:1 |
| Low | Low | High | Lowest | High | High | Mid | 1:1 |
| Low | Low | High | Lowest | High | High | High | 1:1 |
| Mid | Low | Mid | Lowest | n/a | n/a | n/a | n/a |
| Mid | Low | Mid | Lowest | Low | n/a | n/a | n/a |
| Mid | Low | Mid | Lowest | Low | Low | Low | n/a |
| Mid | Low | Mid | Lowest | Low | Mid | Low | n/a |

| $\mu$ | Bottleneck Size | $K$ | Infection Duration | Recombination | $\Xi$ | Burst Size | DfE |
| --- | --- | --- | --- | --- | --- | --- | --- |
| Mid | Low | Mid | Lowest | Low | Low | Low | 4:1 |
| Mid | Low | Mid | Lowest | Low | Low | Mid | 4:1 |
| Mid | Low | Mid | Lowest | Low | Low | High | 4:1 |
| Mid | Low | Mid | Lowest | Low | Mid | Low | 4:1 |
| Mid | Low | Mid | Lowest | Low | Low | Low | 1:1 |
| Mid | Low | Mid | Lowest | Low | Low | Mid | 1:1 |
| Mid | Low | Mid | Lowest | Low | Low | High | 1:1 |
| Mid | Low | Mid | Lowest | Low | Mid | Low | 1:1 |
| Mid | Low | Mid | Lowest | Low | Low | Low | 1:4 |
| Mid | Low | Mid | Lowest | Low | Low | Mid | 1:4 |
| Mid | Low | Mid | Lowest | Low | Mid | Low | 1:4 |
| Mid | Low | Mid | Lowest | Low | High | Low | 1:4 |
| Mid | Low | Mid | Lowest | Mid | n/a | n/a | n/a |
| Mid | Low | Mid | Lowest | Mid | Low | Low | n/a |
| Mid | Low | Mid | Lowest | Mid | Low | Mid | n/a |
| Mid | Low | Mid | Lowest | Mid | Mid | Low | n/a |
| Mid | Low | Mid | Lowest | Mid | Low | Low | 4:1 |
| Mid | Low | Mid | Lowest | Mid | Low | Mid | 4:1 |
| Mid | Low | Mid | Lowest | Mid | Mid | Low | 4:1 |
| Mid | Low | Mid | Lowest | Mid | Low | Low | 1:1 |
| Mid | Low | Mid | Lowest | Mid | Low | Mid | 1:1 |
| Mid | Low | Mid | Lowest | Mid | Low | High | 1:1 |
| Mid | Low | Mid | Lowest | Mid | Mid | Low | 1:1 |
| Mid | Low | Mid | Lowest | Mid | Low | Low | 1:4 |
| Mid | Low | Mid | Lowest | Mid | Low | Mid | 1:4 |
| Mid | Low | Mid | Lowest | Mid | Low | High | 1:4 |
| Mid | Low | Mid | Lowest | Mid | Mid | Low | 1:4 |
| Mid | Low | Mid | Lowest | Mid | Mid | Mid | 1:4 |
| Mid | Low | Mid | Lowest | Mid | High | Low | 1:4 |
| Mid | Low | Mid | Lowest | High | n/a | n/a | n/a |
| Mid | Low | Mid | Lowest | High | Low | Low | n/a |
| Mid | Low | Mid | Lowest | High | Low | Mid | n/a |
| Mid | Low | Mid | Lowest | High | Mid | Low | n/a |
| Mid | Low | Mid | Lowest | High | Low | Low | 4:1 |
| Mid | Low | Mid | Lowest | High | Low | Mid | 4:1 |
| Mid | Low | Mid | Lowest | High | Low | High | 4:1 |
| Mid | Low | Mid | Lowest | High | Mid | Low | 4:1 |
| Mid | Low | Mid | Lowest | High | Low | Low | 1:1 |
| Mid | Low | Mid | Lowest | High | Low | Mid | 1:1 |
| Mid | Low | Mid | Lowest | High | Mid | Low | 1:1 |
| Mid | Low | Mid | Lowest | High | Low | Low | 1:4 |
| Mid | Low | Mid | Lowest | High | Low | Mid | 1:4 |
| Mid | Low | Mid | Lowest | High | Low | High | 1:4 |
| Mid | Low | Mid | Lowest | High | Mid | Low | 1:4 |
| Mid | Low | Mid | Lowest | High | High | Low | 1:4 |
| Mid | Low | High | Lowest | n/a | n/a | n/a | n/a |
| Mid | Low | High | Lowest | Low | n/a | n/a | n/a |
| Mid | Low | High | Lowest | Low | Low | Low | n/a |
| Mid | Low | High | Lowest | Low | Low | Mid | n/a |
| Mid | Low | High | Lowest | Low | Low | High | n/a |
| Mid | Low | High | Lowest | Low | Mid | Low | n/a |
| Mid | Low | High | Lowest | Low | Low | Low | 4:1 |
| Mid | Low | High | Lowest | Low | Low | Mid | 4:1 |
| Mid | Low | High | Lowest | Low | Low | High | 4:1 |
| Mid | Low | High | Lowest | Low | Mid | Low | 4:1 |
| Mid | Low | High | Lowest | Low | High | Low | 4:1 |
| Mid | Low | High | Lowest | Low | Low | Low | 1:1 |
| Mid | Low | High | Lowest | Low | Low | Mid | 1:1 |
| Mid | Low | High | Lowest | Low | Low | High | 1:1 |
| Mid | Low | High | Lowest | Low | Mid | Low | 1:1 |
| Mid | Low | High | Lowest | Low | High | Low | 1:1 |
| Mid | Low | High | Lowest | Low | Low | Low | 1:4 |
| Mid | Low | High | Lowest | Low | Low | Mid | 1:4 |
| Mid | Low | High | Lowest | Low | Low | High | 1:4 |
| Mid | Low | High | Lowest | Low | Mid | Low | 1:4 |
| Mid | Low | High | Lowest | Low | Mid | Mid | 1:4 |
| Mid | Low | High | Lowest | Low | High | Low | 1:4 |
| Mid | Low | High | Lowest | Mid | n/a | n/a | n/a |
| Mid | Low | High | Lowest | Mid | Low | Low | n/a |
| Mid | Low | High | Lowest | Mid | Low | Mid | n/a |
| Mid | Low | High | Lowest | Mid | Mid | Low | n/a |
| Mid | Low | High | Lowest | Mid | Low | Low | 4:1 |
| Mid | Low | High | Lowest | Mid | Low | Mid | 4:1 |
| Mid | Low | High | Lowest | Mid | Low | High | 4:1 |
| Mid | Low | High | Lowest | Mid | Mid | Low | 4:1 |
| Mid | Low | High | Lowest | Mid | Low | Low | 1:1 |
| Mid | Low | High | Lowest | Mid | Low | Mid | 1:1 |
| Mid | Low | High | Lowest | Mid | Low | High | 1:1 |
| Mid | Low | High | Lowest | Mid | Mid | Low | 1:1 |
| Mid | Low | High | Lowest | Mid | Low | Mid | 1:4 |
| Mid | Low | High | Lowest | Mid | Low | High | 1:4 |
| Mid | Low | High | Lowest | Mid | Mid | Low | 1:4 |

| $\mu$ | Bottleneck Size | $K$ | Infection Duration | Recombination | $\Xi$ | Burst Size | DfE |
| --- | --- | --- | --- | --- | --- | --- | --- |
| Mid | Low | High | Lowest | Mid | Mid | Mid | 1:4 |
| Mid | Low | High | Lowest | Mid | High | Mid | 1:4 |
| Mid | Low | High | Lowest | High | n/a | n/a | n/a |
| Mid | Low | High | Lowest | High | Low | Low | n/a |
| Mid | Low | High | Lowest | High | Low | Mid | n/a |
| Mid | Low | High | Lowest | High | Low | High | n/a |
| Mid | Low | High | Lowest | High | Mid | Low | n/a |
| Mid | Low | High | Lowest | High | Low | Low | 4:1 |
| Mid | Low | High | Lowest | High | Low | Mid | 4:1 |
| Mid | Low | High | Lowest | High | Low | High | 4:1 |
| Mid | Low | High | Lowest | High | Mid | Low | 4:1 |
| Mid | Low | High | Lowest | High | Low | Low | 1:1 |
| Mid | Low | High | Lowest | High | Low | Mid | 1:1 |
| Mid | Low | High | Lowest | High | Low | High | 1:1 |
| Mid | Low | High | Lowest | High | Mid | Low | 1:1 |
| Mid | Low | High | Lowest | High | High | Low | 1:1 |
| Mid | Low | High | Lowest | High | Low | Low | 1:4 |
| Mid | Low | High | Lowest | High | Low | Mid | 1:4 |
| Mid | Low | High | Lowest | High | Low | High | 1:4 |
| Mid | Low | High | Lowest | High | Mid | Mid | 1:4 |
| Mid | Low | High | Lowest | High | High | Low | 1:4 |
| High | Low | High | Lowest | Low | Low | Low | 1:4 |
| High | Low | High | Lowest | Mid | Low | Low | 1:4 |
| High | Low | High | Lowest | High | Low | Mid | 1:4 |
| Low | Low | Low | Low | Low | Mid | High | 1:1 |
| Low | Low | Low | Low | Low | Low | Low | 1:4 |
| Low | Low | Low | Low | Low | Low | Mid | 1:4 |
| Low | Low | Low | Low | Low | Low | High | 1:4 |
| Low | Low | Low | Low | Low | Mid | Low | 1:4 |
| Low | Low | Low | Low | Low | Mid | Mid | 1:4 |
| Low | Low | Low | Low | Low | Mid | High | 1:4 |
| Low | Low | Low | Low | Low | High | Low | 1:4 |
| Low | Low | Low | Low | Low | High | Mid | 1:4 |
| Low | Low | Low | Low | Low | High | High | 1:4 |
| Low | Low | Low | Low | Mid | Mid | High | 1:1 |
| Low | Low | Low | Low | Mid | Low | Low | 1:4 |
| Low | Low | Low | Low | Mid | Low | Mid | 1:4 |
| Low | Low | Low | Low | Mid | Low | High | 1:4 |
| Low | Low | Low | Low | Mid | Mid | Low | 1:4 |
| Low | Low | Low | Low | Mid | Mid | Mid | 1:4 |
| Low | Low | Low | Low | Mid | Mid | High | 1:4 |
| Low | Low | Low | Low | Mid | High | Low | 1:4 |
| Low | Low | Low | Low | Mid | High | Mid | 1:4 |
| Low | Low | Low | Low | Mid | High | High | 1:4 |
| Low | Low | Low | Low | High | Mid | High | 1:1 |
| Low | Low | Low | Low | High | Low | Low | 1:4 |
| Low | Low | Low | Low | High | Low | Mid | 1:4 |
| Low | Low | Low | Low | High | Low | High | 1:4 |
| Low | Low | Low | Low | High | Mid | Low | 1:4 |
| Low | Low | Low | Low | High | Mid | High | 1:4 |
| Low | Low | Low | Low | High | High | Low | 1:4 |
| Low | Low | Low | Low | High | High | High | 1:4 |
| Low | Low | Mid | Low | n/a | n/a | n/a | n/a |
| Low | Low | Mid | Low | Low | n/a | n/a | n/a |
| Low | Low | Mid | Low | Low | Low | Low | n/a |
| Low | Low | Mid | Low | Low | Low | Mid | n/a |
| Low | Low | Mid | Low | Low | Low | High | n/a |
| Low | Low | Mid | Low | Low | Mid | Low | n/a |
| Low | Low | Mid | Low | Low | Low | Mid | 4:1 |
| Low | Low | Mid | Low | Low | Low | High | 4:1 |
| Low | Low | Mid | Low | Low | Mid | Low | 4:1 |
| Low | Low | Mid | Low | Low | Low | Mid | 1:1 |
| Low | Low | Mid | Low | Low | Low | High | 1:1 |
| Low | Low | Mid | Low | Low | Mid | Low | 1:1 |
| Low | Low | Mid | Low | Low | High | Low | 1:1 |
| Low | Low | Mid | Low | Low | Low | High | 1:4 |
| Low | Low | Mid | Low | Low | Mid | Mid | 1:4 |
| Low | Low | Mid | Low | Low | Mid | High | 1:4 |
| Low | Low | Mid | Low | Low | High | Low | 1:4 |
| Low | Low | Mid | Low | Mid | n/a | n/a | n/a |
| Low | Low | Mid | Low | Mid | Low | Low | n/a |
| Low | Low | Mid | Low | Mid | Low | Mid | n/a |
| Low | Low | Mid | Low | Mid | Low | High | n/a |
| Low | Low | Mid | Low | Mid | Low | Low | 4:1 |
| Low | Low | Mid | Low | Mid | Low | Mid | 4:1 |
| Low | Low | Mid | Low | Mid | Low | High | 4:1 |
| Low | Low | Mid | Low | Mid | Mid | Low | 4:1 |
| Low | Low | Mid | Low | Mid | High | Low | 1:1 |
| Low | Low | Mid | Low | Mid | Low | High | 1:4 |
| Low | Low | Mid | Low | Mid | Mid | Mid | 1:4 |
| Low | Low | Mid | Low | Mid | Mid | High | 1:4 |
| Low | Low | Mid | Low | Mid | High | Low | 1:4 |
| Low | Low | Mid | Low | Mid | High | Mid | 1:4 |
| Low | Low | Mid | Low | Mid | High | High | 1:4 |
| Low | Low | Mid | Low | High | n/a | n/a | n/a |
| Low | Low | Mid | Low | High | Low | Low | n/a |
| Low | Low | Mid | Low | High | Low | Mid | n/a |
| Low | Low | Mid | Low | High | Low | High | n/a |
| Low | Low | Mid | Low | High | Low | Mid | 4:1 |
| Low | Low | Mid | Low | High | Low | High | 4:1 |
| Low | Low | Mid | Low | High | Mid | Low | 4:1 |
| Low | Low | Mid | Low | High | Low | Low | 1:1 |
| Low | Low | Mid | Low | High | Mid | Low | 1:1 |
| Low | Low | Mid | Low | High | Mid | Mid | 1:1 |
| Low | Low | Mid | Low | High | Mid | Mid | 1:4 |
| Low | Low | Mid | Low | High | Mid | High | 1:4 |
| Low | Low | Mid | Low | High | High | Low | 1:4 |
| Low | Low | Mid | Low | High | High | Mid | 1:4 |
| Low | Low | Mid | Low | High | High | High | 1:4 |
| Low | Low | High | Low | Low | n/a | n/a | n/a |
| Low | Low | High | Low | Low | Low | Low | n/a |
| Low | Low | High | Low | Low | Low | Mid | n/a |
| Low | Low | High | Low | Low | Mid | Low | n/a |
| Low | Low | High | Low | Low | High | Low | n/a |
| Low | Low | High | Low | Low | Low | Low | 4:1 |
| Low | Low | High | Low | Low | Mid | Mid | 4:1 |
| Low | Low | High | Low | Low | High | Low | 4:1 |
| Low | Low | High | Low | Low | Low | High | 1:1 |
| Low | Low | High | Low | Low | Mid | Low | 1:1 |
| Low | Low | High | Low | Low | Mid | Mid | 1:1 |
| Low | Low | High | Low | Low | High | Low | 1:1 |
| Low | Low | High | Low | Low | Mid | Mid | 1:4 |
| Low | Low | High | Low | Low | Mid | High | 1:4 |
| Low | Low | High | Low | Low | High | Low | 1:4 |
| Low | Low | High | Low | Low | High | Mid | 1:4 |
| Low | Low | High | Low | Low | High | High | 1:4 |
| Low | Low | High | Low | Mid | n/a | n/a | n/a |
| Low | Low | High | Low | Mid | Low | Low | n/a |
| Low | Low | High | Low | Mid | Low | High | n/a |
| Low | Low | High | Low | Mid | Mid | Low | n/a |
| Low | Low | High | Low | Mid | High | Low | n/a |
| Low | Low | High | Low | Mid | Low | Low | 4:1 |
| Low | Low | High | Low | Mid | Low | Mid | 4:1 |
| Low | Low | High | Low | Mid | Low | High | 4:1 |
| Low | Low | High | Low | Mid | Mid | Low | 4:1 |
| Low | Low | High | Low | Mid | Mid | Mid | 4:1 |
| Low | Low | High | Low | Mid | High | Low | 4:1 |
| Low | Low | High | Low | Mid | Low | Mid | 1:1 |
| Low | Low | High | Low | Mid | Mid | Low | 1:1 |
| Low | Low | High | Low | Mid | Mid | Mid | 1:1 |
| Low | Low | High | Low | Mid | High | Low | 1:1 |
| Low | Low | High | Low | Mid | High | Mid | 1:1 |
| Low | Low | High | Low | Mid | Low | Low | 1:4 |
| Low | Low | High | Low | Mid | Mid | Mid | 1:4 |
| Low | Low | High | Low | Mid | Mid | High | 1:4 |
| Low | Low | High | Low | Mid | High | Mid | 1:4 |
| Low | Low | High | Low | Mid | High | High | 1:4 |
| Low | Low | High | Low | High | n/a | n/a | n/a |
| Low | Low | High | Low | High | Low | Low | n/a |
| Low | Low | High | Low | High | Low | Mid | n/a |
| Low | Low | High | Low | High | Mid | Low | n/a |
| Low | Low | High | Low | High | High | Low | n/a |
| Low | Low | High | Low | High | Low | Mid | 4:1 |
| Low | Low | High | Low | High | Mid | Mid | 4:1 |
| Low | Low | High | Low | High | High | Low | 4:1 |
| Low | Low | High | Low | High | Low | Low | 1:1 |
| Low | Low | High | Low | High | Low | Mid | 1:1 |
| Low | Low | High | Low | High | Mid | Mid | 1:1 |
| Low | Low | High | Low | High | High | High | 1:1 |
| Low | Low | High | Low | High | Low | High | 1:4 |
| Low | Low | High | Low | High | Mid | Mid | 1:4 |
| Low | Low | High | Low | High | Mid | High | 1:4 |
| Low | Low | High | Low | High | High | Mid | 1:4 |
| Low | Low | High | Low | High | High | High | 1:4 |
| Mid | Low | Mid | Low | Low | Low | Low | 1:4 |
| Mid | Low | Mid | Low | Low | Low | Mid | 1:4 |
| Mid | Low | Mid | Low | Low | Mid | Low | 1:4 |
| Mid | Low | Mid | Low | Mid | Low | Low | 1:4 |
| Mid | Low | Mid | Low | Mid | Low | Mid | 1:4 |
| Mid | Low | Mid | Low | Mid | Mid | Low | 1:4 |
| Mid | Low | Mid | Low | High | Low | Low | 1:4 |
| Mid | Low | Mid | Low | High | Low | Mid | 1:4 |
| Mid | Low | Mid | Low | High | Mid | Low | 1:4 |
| Mid | Low | High | Low | Low | n/a | n/a | n/a |
| Mid | Low | High | Low | Low | Low | Low | n/a |
| Mid | Low | High | Low | Low | Low | Mid | n/a |
| Mid | Low | High | Low | Low | Low | Low | 4:1 |
| Mid | Low | High | Low | Low | Low | Low | 1:1 |
| Mid | Low | High | Low | Low | Low | Low | 1:4 |
| Mid | Low | High | Low | Mid | n/a | n/a | n/a |
| Mid | Low | High | Low | Mid | Low | Low | n/a |
| Mid | Low | High | Low | Mid | Low | Low | 4:1 |
| Mid | Low | High | Low | Mid | Low | Mid | 4:1 |
| Mid | Low | High | Low | Mid | Low | Low | 1:1 |
| Mid | Low | High | Low | Mid | Low | Mid | 1:1 |
| Mid | Low | High | Low | Mid | Mid | Low | 1:1 |
| Mid | Low | High | Low | Mid | Low | Low | 1:4 |
| Mid | Low | High | Low | Mid | Low | Mid | 1:4 |
| Mid | Low | High | Low | Mid | Low | High | 1:4 |
| Mid | Low | High | Low | Mid | Mid | Low | 1:4 |
| Mid | Low | High | Low | High | n/a | n/a | n/a |
| Mid | Low | High | Low | High | Low | Low | 4:1 |
| Mid | Low | High | Low | High | Low | Low | 1:1 |
| Mid | Low | High | Low | High | Low | Mid | 1:1 |
| Mid | Low | High | Low | High | Mid | Low | 1:1 |
| Mid | Low | High | Low | High | Low | Low | 1:4 |
| Mid | Low | High | Low | High | Low | Mid | 1:4 |
| Mid | Low | High | Low | High | Low | High | 1:4 |
| Mid | Low | High | Low | High | Mid | Low | 1:4 |
| Mid | Low | High | Low | High | High | Low | 1:4 |
| Low | Low | Low | Mid | Low | Mid | High | 1:1 |
| Low | Low | Low | Mid | Low | Mid | Mid | 1:4 |
| Low | Low | Low | Mid | Low | Mid | High | 1:4 |
| Low | Low | Low | Mid | Mid | Mid | High | 1:1 |
| Low | Low | Low | Mid | Mid | Mid | High | 1:4 |
| Low | Low | Low | Mid | High | Mid | High | 1:1 |
| Low | Low | Low | Mid | High | Low | High | 1:4 |
| Low | Low | Low | Mid | High | Mid | High | 1:4 |
| Low | Low | Low | Mid | High | High | Mid | 1:4 |
| Low | Low | Mid | Mid | n/a | n/a | n/a | n/a |
| Low | Low | Mid | Mid | Low | n/a | n/a | n/a |
| Low | Low | Mid | Mid | Low | Low | Low | n/a |
| Low | Low | Mid | Mid | Low | Low | Low | 4:1 |
| Low | Low | Mid | Mid | Low | Mid | Low | 4:1 |
| Low | Low | Mid | Mid | Low | Low | Low | 1:1 |
| Low | Low | Mid | Mid | Low | Low | Mid | 1:1 |
| Low | Low | Mid | Mid | Low | Low | High | 1:1 |
| Low | Low | Mid | Mid | Low | Mid | Low | 1:1 |
| Low | Low | Mid | Mid | Low | Low | Low | 1:4 |
| Low | Low | Mid | Mid | Low | Low | Mid | 1:4 |
| Low | Low | Mid | Mid | Low | Low | High | 1:4 |
| Low | Low | Mid | Mid | Low | Mid | Low | 1:4 |
| Low | Low | Mid | Mid | Mid | Low | Low | 4:1 |
| Low | Low | Mid | Mid | Mid | Mid | Low | 4:1 |
| Low | Low | Mid | Mid | Mid | Low | Low | 1:1 |
| Low | Low | Mid | Mid | Mid | Low | Mid | 1:1 |
| Low | Low | Mid | Mid | Mid | Mid | Low | 1:1 |
| Low | Low | Mid | Mid | Mid | Low | Mid | 1:4 |
| Low | Low | Mid | Mid | Mid | Low | High | 1:4 |
| Low | Low | Mid | Mid | Mid | Mid | Low | 1:4 |
| Low | Low | Mid | Mid | High | n/a | n/a | n/a |
| Low | Low | Mid | Mid | High | Low | Mid | 1:1 |
| Low | Low | Mid | Mid | High | Low | Low | 1:4 |
| Low | Low | Mid | Mid | High | Low | Mid | 1:4 |
| Low | Low | Mid | Mid | High | Low | High | 1:4 |
| Low | Low | Mid | Mid | High | Mid | Low | 1:4 |
| Low | Low | High | Mid | n/a | n/a | n/a | n/a |
| Low | Low | High | Mid | Low | n/a | n/a | n/a |
| Low | Low | High | Mid | Low | Low | Mid | n/a |
| Low | Low | High | Mid | Low | Mid | Low | n/a |
| Low | Low | High | Mid | Low | Low | Mid | 4:1 |
| Low | Low | High | Mid | Low | Low | High | 4:1 |
| Low | Low | High | Mid | Low | Low | Low | 1:1 |
| Low | Low | High | Mid | Low | Low | Mid | 1:1 |
| Low | Low | High | Mid | Low | Low | High | 1:1 |
| Low | Low | High | Mid | Low | Mid | Low | 1:1 |
| Low | Low | High | Mid | Low | Low | Mid | 1:4 |
| Low | Low | High | Mid | Low | Mid | Mid | 1:4 |
| Low | Low | High | Mid | Low | High | Low | 1:4 |
| Low | Low | High | Mid | Mid | n/a | n/a | n/a |
| Low | Low | High | Mid | Mid | Low | Low | n/a |
| Low | Low | High | Mid | Mid | Low | Mid | n/a |
| Low | Low | High | Mid | Mid | Mid | Low | n/a |
| Low | Low | High | Mid | Mid | Low | Low | 4:1 |
| Low | Low | High | Mid | Mid | Low | Low | 1:1 |
| Low | Low | High | Mid | Mid | Low | Mid | 1:1 |
| Low | Low | High | Mid | Mid | Low | High | 1:4 |
| Low | Low | High | Mid | Mid | Mid | Mid | 1:4 |
| Low | Low | High | Mid | Mid | High | Low | 1:4 |
| Low | Low | High | Mid | High | n/a | n/a | n/a |
| Low | Low | High | Mid | High | Low | Low | n/a |
| Low | Low | High | Mid | High | Low | Mid | n/a |
| Low | Low | High | Mid | High | Low | High | n/a |
| Low | Low | High | Mid | High | Mid | Low | n/a |
| Low | Low | High | Mid | High | Low | Mid | 4:1 |
| Low | Low | High | Mid | High | Low | High | 4:1 |
| Low | Low | High | Mid | High | Mid | Low | 4:1 |
| Low | Low | High | Mid | High | Low | Low | 1:1 |
| Low | Low | High | Mid | High | Low | High | 1:1 |
| Low | Low | High | Mid | High | High | Low | 1:1 |
| Low | Low | High | Mid | High | Low | High | 1:4 |
| Low | Low | High | Mid | High | Mid | Low | 1:4 |
| Low | Low | High | Mid | High | Mid | Mid | 1:4 |
| Low | Low | High | Mid | High | High | Low | 1:4 |
| Mid | Low | High | Mid | Low | Low | Mid | 1:4 |
| Mid | Low | High | Mid | Mid | Low | Mid | 1:4 |
| Mid | Low | High | Mid | High | Low | Mid | 1:4 |
| Low | Low | Low | High | Low | Mid | Mid | 1:4 |
| Low | Low | Low | High | Low | Mid | High | 1:4 |
| Low | Low | Low | High | Mid | Mid | Mid | 1:4 |
| Low | Low | Low | High | Mid | Mid | High | 1:4 |
| Low | Low | Low | High | High | Mid | High | 1:4 |
| Low | Low | Mid | High | Low | Low | Low | 1:4 |
| Low | Low | Mid | High | Low | Low | Mid | 1:4 |
| Low | Low | Mid | High | Low | Mid | Low | 1:4 |
| Low | Low | Mid | High | Mid | Low | Low | 1:4 |
| Low | Low | Mid | High | Mid | Low | Mid | 1:4 |
| Low | Low | Mid | High | Mid | Low | High | 1:4 |
| Low | Low | Mid | High | Mid | Mid | Low | 1:4 |
| Low | Low | Mid | High | High | Low | Low | 1:4 |
| Low | Low | Mid | High | High | Low | Mid | 1:4 |
| Low | Low | Mid | High | High | Mid | Low | 1:4 |
| Low | Low | High | High | n/a | n/a | n/a | n/a |
| Low | Low | High | High | Low | Low | Low | 4:1 |
| Low | Low | High | High | Low | Low | Mid | 4:1 |
| Low | Low | High | High | Low | Low | Low | 1:1 |
| Low | Low | High | High | Low | Low | Mid | 1:1 |

| $\mu$ | Bottleneck<br>Size | $K$ | Infection<br>Duration | Recombination | $\Xi$ | Burst<br>Size | DFE |
| --- | --- | --- | --- | --- | --- | --- | --- |
| Low | Low | High | High | Low | Mid | Low | 1:1 |
| Low | Low | High | High | Low | Low | Mid | 1:4 |
| Low | Low | High | High | Low | Low | High | 1:4 |
| Low | Low | High | High | Low | Mid | Low | 1:4 |
| Low | Low | High | High | Mid | Low | Low | 4:1 |
| Low | Low | High | High | Mid | Low | Low | 1:1 |
| Low | Low | High | High | Mid | Low | Mid | 1:1 |
| Low | Low | High | High | Mid | Mid | Low | 1:1 |
| Low | Low | High | High | Mid | Low | Low | 1:4 |
| Low | Low | High | High | Mid | Low | Mid | 1:4 |
| Low | Low | High | High | Mid | Low | High | 1:4 |
| Low | Low | High | High | Mid | Mid | Low | 1:4 |
| Low | Low | High | High | High | n/a | n/a | n/a |
| Low | Low | High | High | High | Low | Low | 4:1 |
| Low | Low | High | High | High | Low | Mid | 1:1 |
| Low | Low | High | High | High | Mid | Low | 1:1 |
| Low | Low | High | High | High | Low | Low | 1:4 |
| Low | Low | High | High | High | Low | Mid | 1:4 |
| Low | Low | High | High | High | Low | High | 1:4 |
| Low | Low | High | High | High | Mid | Low | 1:4 |

**Supplemental Table 2.**
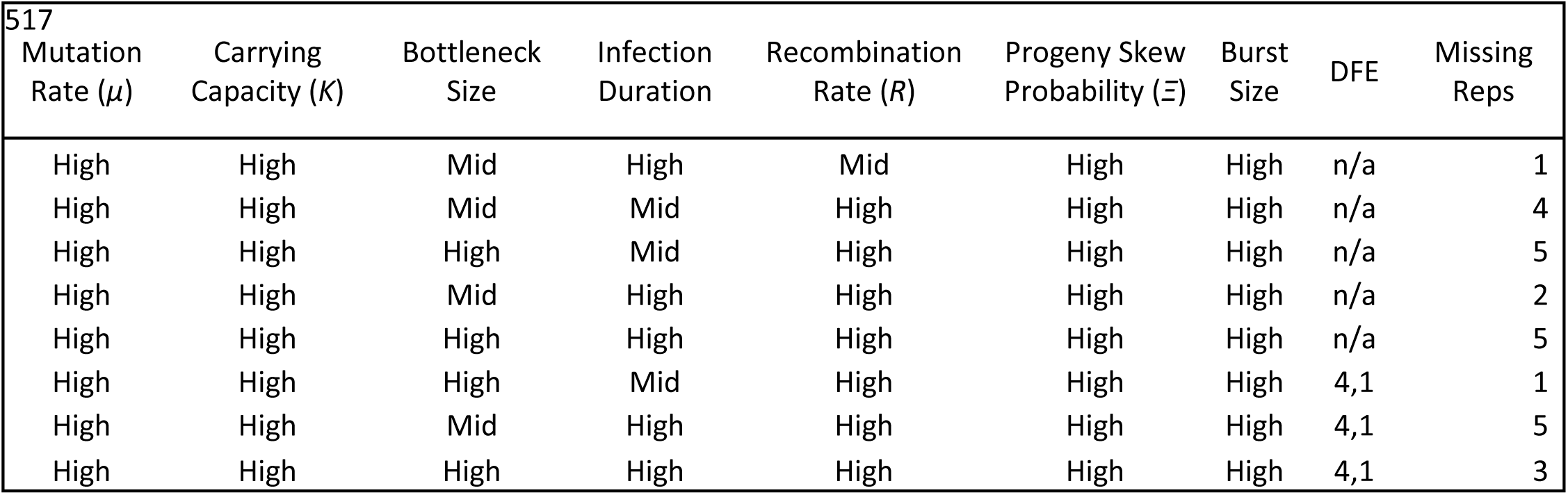
List of models with replicates that failed to complete due to exceeding allocated memory.

**Supplemental Table 3.** Order of models presented in Figure 3, Supplemental Figure 1, and Supplemental Figure 2 as lines (range) and points (mean value). Note that for each figure the mutation rate (*µ*), initial bottleneck size, and carrying capacity (*K*) are the same for all models. The infection duration varies between models as described in figure captions. Models with no completed replicates (Supplemental Table 2) appear as gaps between neighboring models.

| Order | $R$ | $\Xi$ | Burst Size | DFE |
| --- | --- | --- | --- | --- |
| 1 | n/a | n/a | n/a | n/a |
| 2 | Low | n/a | n/a | n/a |
| 3 | Mid | n/a | n/a | n/a |
| 4 | High | n/a | n/a | n/a |
| 5 | Low | Low | Low | n/a |
| 6 | Mid | Low | Low | n/a |
| 7 | High | Low | Low | n/a |
| 8 | Low | Mid | Low | n/a |
| 9 | Mid | Mid | Low | n/a |
| 10 | High | Mid | Low | n/a |
| 11 | Low | High | Low | n/a |
| 12 | Mid | High | Low | n/a |
| 13 | High | High | Low | n/a |
| 14 | Low | Low | Mid | n/a |
| 15 | Mid | Low | Mid | n/a |
| 16 | High | Low | Mid | n/a |
| 17 | Low | Mid | Mid | n/a |
| 18 | Mid | Mid | Mid | n/a |
| 19 | High | Mid | Mid | n/a |
| 20 | Low | High | Mid | n/a |
| 21 | Mid | High | Mid | n/a |
| 22 | High | High | Mid | n/a |
| 23 | Low | Low | High | n/a |
| 24 | Mid | Low | High | n/a |
| 25 | High | Low | High | n/a |
| 26 | Low | Mid | High | n/a |
| 27 | Mid | Mid | High | n/a |
| 28 | High | Mid | High | n/a |
| 29 | Low | High | High | n/a |
| 30 | Mid | High | High | n/a |
| 31 | High | High | High | n/a |
| 32 | Low | Low | Low | 4:1 |
| 33 | Mid | Low | Low | 4:1 |
| 34 | High | Low | Low | 4:1 |
| 35 | Low | Mid | Low | 4:1 |
| 36 | Mid | Mid | Low | 4:1 |
| 37 | High | Mid | Low | 4:1 |
| 38 | Low | High | Low | 4:1 |
| 39 | Mid | High | Low | 4:1 |
| 40 | High | High | Low | 4:1 |
| 41 | Low | Low | Mid | 4:1 |
| 42 | Mid | Low | Mid | 4:1 |
| 43 | High | Low | Mid | 4:1 |
| 44 | Low | Mid | Mid | 4:1 |
| 45 | Mid | Mid | Mid | 4:1 |
| 46 | High | Mid | Mid | 4:1 |
| 47 | Low | High | Mid | 4:1 |
| 48 | Mid | High | Mid | 4:1 |
| 49 | High | High | Mid | 4:1 |
| 50 | Low | Low | High | 4:1 |
| 51 | Mid | Low | High | 4:1 |
| 52 | High | Low | High | 4:1 |
| 53 | Low | Mid | High | 4:1 |
| 54 | Mid | Mid | High | 4:1 |
| 55 | High | Mid | High | 4:1 |
| 56 | Low | High | High | 4:1 |
| 57 | Mid | High | High | 4:1 |
| 58 | High | High | High | 4:1 |
| 59 | Low | Low | Low | 1:1 |
| 60 | Mid | Low | Low | 1:1 |
| 61 | High | Low | Low | 1:1 |
| 62 | Low | Mid | Low | 1:1 |
| 63 | Mid | Mid | Low | 1:1 |
| 64 | High | Mid | Low | 1:1 |
| 65 | Low | High | Low | 1:1 |
| 66 | Mid | High | Low | 1:1 |
| 67 | High | High | Low | 1:1 |
| 68 | Low | Low | Mid | 1:1 |
| 69 | Mid | Low | Mid | 1:1 |
| 70 | High | Low | Mid | 1:1 |
| 71 | Low | Mid | Mid | 1:1 |
| 72 | Mid | Mid | Mid | 1:1 |
| 73 | High | Mid | Mid | 1:1 |
| 74 | Low | High | Mid | 1:1 |
| 75 | Mid | High | Mid | 1:1 |
| 76 | High | High | Mid | 1:1 |
| 77 | Low | Low | High | 1:1 |
| 78 | Mid | Low | High | 1:1 |
| 79 | High | Low | High | 1:1 |
| 80 | Low | Mid | High | 1:1 |
| 81 | Mid | Mid | High | 1:1 |
| 82 | High | Mid | High | 1:1 |
| 83 | Low | High | High | 1:1 |
| 84 | Mid | High | High | 1:1 |
| 85 | High | High | High | 1:1 |
| 86 | Low | Low | Low | 1:4 |
| 87 | Mid | Low | Low | 1:4 |
| 88 | High | Low | Low | 1:4 |
| 89 | Low | Mid | Low | 1:4 |
| 90 | Mid | Mid | Low | 1:4 |
| 91 | High | Mid | Low | 1:4 |
| 92 | Low | High | Low | 1:4 |
| 93 | Mid | High | Low | 1:4 |
| 94 | High | High | Low | 1:4 |
| 95 | Low | Low | Mid | 1:4 |
| 96 | Mid | Low | Mid | 1:4 |
| 97 | High | Low | Mid | 1:4 |
| 98 | Low | Mid | Mid | 1:4 |
| 99 | Mid | Mid | Mid | 1:4 |
| 100 | High | Mid | Mid | 1:4 |
| 101 | Low | High | Mid | 1:4 |
| 102 | Mid | High | Mid | 1:4 |
| 103 | High | High | Mid | 1:4 |
| 104 | Low | Low | High | 1:4 |
| 105 | Mid | Low | High | 1:4 |
| 106 | High | Low | High | 1:4 |
| 107 | Low | Mid | High | 1:4 |
| 108 | Mid | Mid | High | 1:4 |
| 109 | High | Mid | High | 1:4 |
| 110 | Low | High | High | 1:4 |
| 111 | Mid | High | High | 1:4 |
| 112 | High | High | High | 1:4 |

**Supplemental Figure 1.**
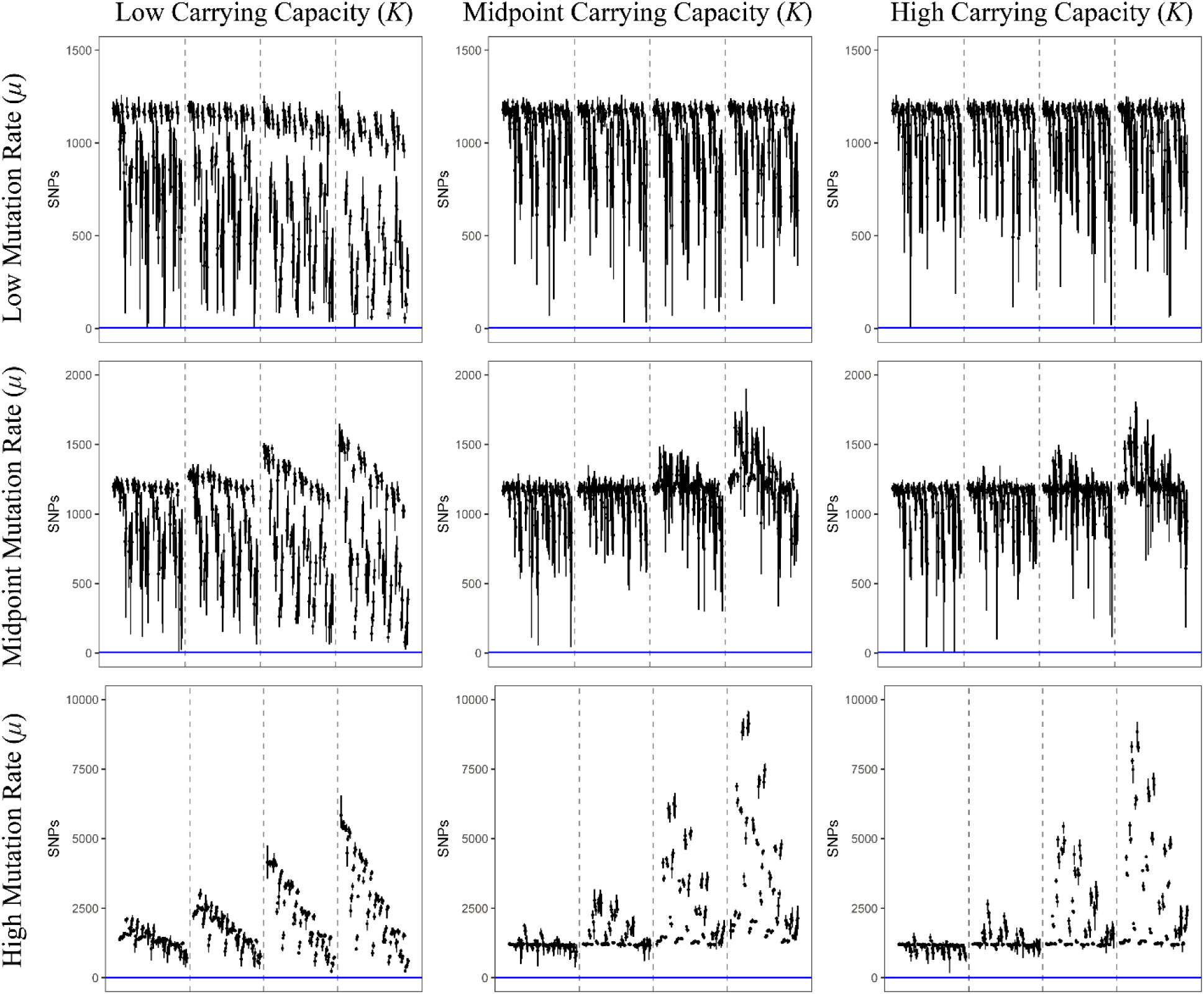
Each line represents the range of filtered SNPs for a particular model using a sampling of 1000 genomes; each point is the mean of that model’s replicates. All models in this figure used the lowest bottleneck size (*i.e*., 5); panels increase from left to right in the value of carrying capacity used and increase from top to bottom in the value of mutation rate used. Within each panel, dashed lines separate models into subpanels with different infection durations, increasing from left to right. Within each subpanel the order of the models is the same and is detailed in Supplemental Table 3. The blue, horizontal line represents the threshold of 5 SNPs used in this study to accept or reject a potential model (note that Y-axes differ by row).

**Supplemental Figure 2.**
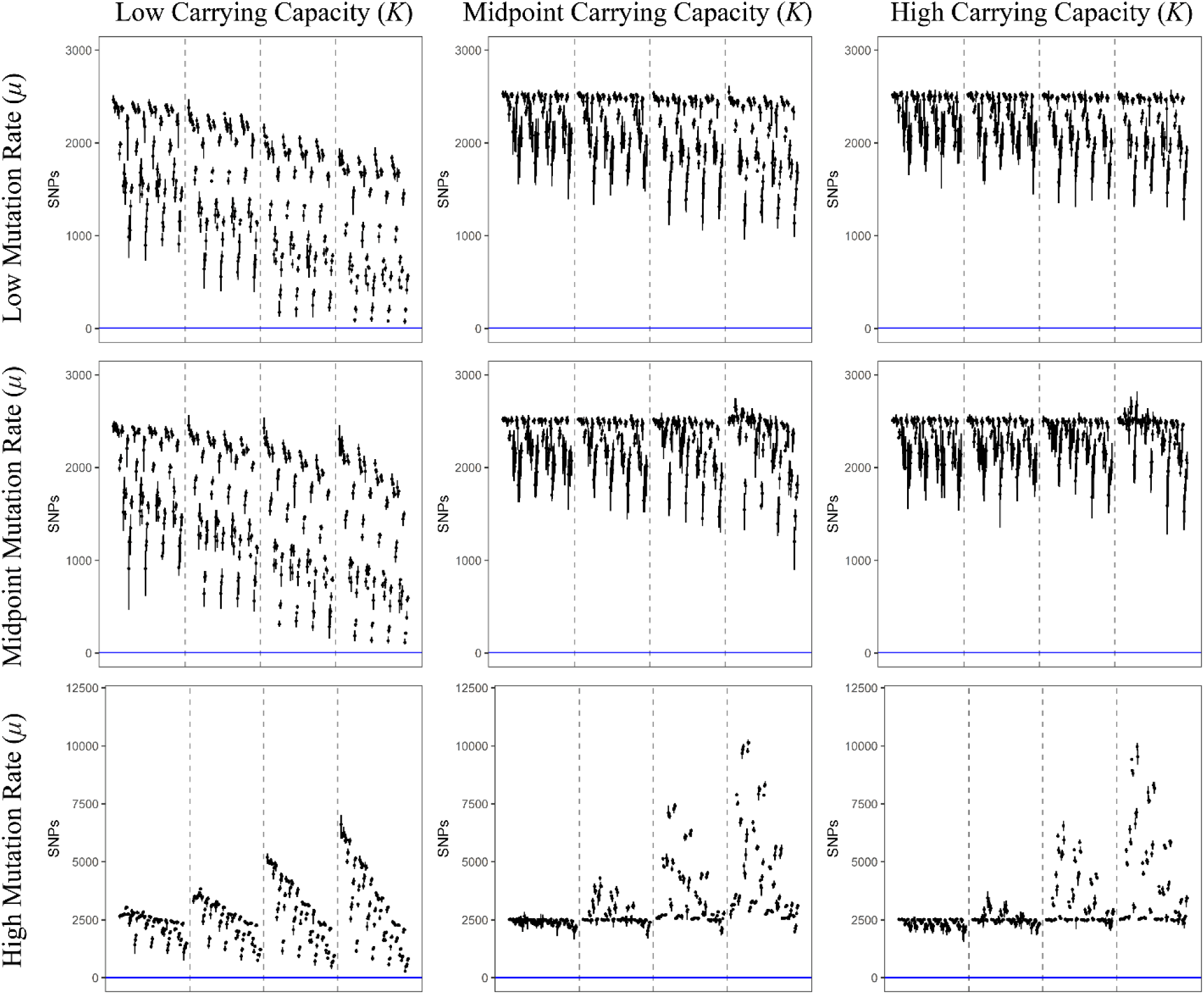
Each line represents the range of filtered SNPs for a particular model using a sampling of 1000 genomes; each point is the mean of that model’s replicates. All models in this figure used the high bottleneck size (*i.e*., 100); panels increase from left to right in the value of carrying capacity used and increase from top to bottom in the value of mutation rate used. Within each panel, dashed lines separate models into subpanels with different infection durations, increasing from left to right. Within each subpanel the order of the models is the same and is detailed in Supplemental Table 3. The blue, horizontal line represents the threshold of 5 SNPs used in this study to accept or reject a potential model (note that Y-axes differ by row).

